# The Endosomal Sorting Complex, ESCRT, has diverse roles in blood progenitor maintenance, lineage choice and immune response

**DOI:** 10.1101/2021.11.29.470366

**Authors:** Arindam Ray, Yashashwinee Rai, Maneesha S Inamdar

## Abstract

Most hematological malignancies are associated with reduced expression of one or more components of the Endosomal Sorting Complex Required for Transport (ESCRT). However, the roles of ESCRT in stem cell and progenitor maintenance are not resolved. The difficulty in parsing signaling pathway roles in relation to their canonical cargo sorting function poses a challenge. The *Drosophila* hematopoietic organ, the larval lymph gland, provides a path to dissect the roles of cellular trafficking pathways such as ESCRT in blood development and maintenance. *Drosophila* has 13 core ESCRT components. Knockdown of individual ESCRTs showed that only Vps28 and Vp36 were required in all lymph gland progenitors. Using the well-conserved ESCRT-II complex (Vps22, Vps25 and Vps36) as an example of the range of phenotypes seen upon ESCRT depletion, we show that ESCRTs have cell autonomous as well as non-autonomous roles in progenitor maintenance and differentiation. ESCRT depletion also sensitized posterior lymph gland progenitors to respond to immunogenic cues such as wasp infestation. We also identify key heterotypic roles for ESCRT in position-dependent control of Notch activation to suppress crystal cell differentiation. Our study shows that the cargo sorting machinery can determine progenitor identity and capacity to adapt to the dynamic environments that blood cells are exposed to. These mechanisms for control of cell fate may tailor developmental diversity in multiple contexts.

## Introduction

Tissue patterning requires spatiotemporally controlled cell proliferation, progenitor specification and lineage differentiation. While a limited number of signaling circuits impact these complex cell properties, their regulation is highly context-dependent. Endocytic trafficking integrates extracellular cues with intracellular changes to maintain, attenuate or amplify signaling [1]. The stereotypical roles of the endocytic machinery in transport and cargo sorting converge to generate complex signaling, thereby significantly impacting tissue homeostasis. However, due to their overlapping roles, tissue and cell type-specific functions of the sorting machinery components have been difficult to dissect out in vivo.

Protein trafficking and turnover through the endo-lysosomal route allows rapid post-translational modulation of signal transduction. The conserved ESCRT machinery actively controls the sorting of ubiquitinated cargoes for lysosomal degradation. This complex consists of four hetero-oligomeric subunits (ESCRT-0, I, II and III) that sequentially bind to endomembrane-bound ubiquitinated cargoes in order to sequester them into intraluminal vesicles (ILV) of the multivesicular bodies/endosomes (MVB/MVE). The *Drosophila* ESCRT is comprised of 13 core components [2, 3]. ESCRT-0 (Hrs, Stam) binds to the ubiquitinated cargoes through a ubiquitin-interacting motif. It then recruits ESCRT-I (Vps28, Tsg101, Vps37A, Vps37B) and ESCRT-II (Vps25, Vps22 and Vps36), which act as a bridging complex to assemble ESCRT-III (Vps32, Vps24, Vps20, Vps2). ESCRT-I-dependent membrane inward budding and ESCRT-III-dependent membrane scission lie at the heart of endosomal protein sorting [4]. Vps32 is the principal filament-forming component that undergoes activation and polymerization upon binding with various nucleating factors and integrates previous steps of endosomal sorting [5]. The final step involves disassembly of ESCRT subunits and scission of the membrane neck of the intraluminal vesicles, which is mediated by the Vps4-Vta1 mechanoenzyme complex.

Though phenotypic diversity of ESCRT mutants is rare in unicellular organisms like budding yeast, dysfunction of metazoan ESCRT components can manifest as distinct and diverse cellular and histological phenotypes such as defective MVB biogenesis and incorrect cell fate choice, tissue hyperproliferation, apoptotic resistance, neoplastic transformation etc., due to dysregulated activation of signaling pathways such as Notch, EGFR and JAK/STAT, thereby altering tissue homeostasis [2, 5-7]. The range of functional outputs of ESCRT modulation makes it an interesting target to explore in the context of stem and progenitor cell lineage choice.

While several endocytic proteins such as Atg6 [8], WASH [9], Rabex-5 [10], Rab5, Rab 11 [11], Asrij [12] and ARF-1 [13] are implicated in developmental signaling and blood cell homeostasis, little is known about the role of ESCRT in hematopoiesis. Previous genetic screens and knockout-based functional analyses in both *Drosophila* and mouse models showed a role of ESCRT in maturation of specific blood cell types in erythroid and lymphoid lineages and a possible functional link of ESCRT to blood cell homeostasis [14-16]. Hence, we analysed the role of ESCRT in blood cell homeostasis, using the *Drosophila* lymph gland model.

The larval blood progenitors in *Drosophila* reside in the multi-lobed lymph gland that flanks the cardiac tube in segments T3 and A1. The primary lobe has a medullary zone enriched in progenitors, cortical zone of differentiated blood cells (plasmatocytes, crystal cells and lamellocytes) and the hematopoietic niche (posterior signaling centre). Blood progenitors of *Drosophila* are linearly arranged in primary, secondary and tertiary lobes of the lymph gland and are characterized by the expression of several markers such as Domeless, TepIV and DE-Cadherin. The lymph gland develops in an anterior to posterior sequence, with younger progenitors in the posterior lobes [17], allowing complete sampling of the progenitor pool. Previous studies showed heterogeneity of the progenitor population in gene expression, mitochondrial morphology and dynamics, signaling, differentiation potential and immune function [17, 18]. Posterior progenitors are refractile to immune challenge due to differential activation of JAK/STAT and Notch signaling [17, 18]. Thus, the lymph gland is an accessible model representing the complexities of tissue homeostasis. Hence, we investigated the role of ESCRT components in spatiotemporal control of blood progenitor homeostasis and myelopoiesis in the lymph gland.

Here, we elucidate the role of all 13 *Drosophila* core ESCRT components in ubiquitinated cargo sorting and blood cell lineage choice across progenitor subsets. We show that though ubiquitous, each ESCRT can have progenitor-specific and lineage-restricted effects mediated by differential regulation of intra- and intercellular signaling. Further, we find that ESCRT perturbation affects Notch mediated crystal cell differentiation. Our study provides a means to deconstruct the roles of ESCRT in spatiotemporal segregation of signaling, giving further insight into progenitor heterogeneity.

## Results

### ESCRT components are mis-expressed in hematological malignancies

The **M**icroarray **I**nnovations in **Le**ukaemia (MILE) study is a collection of gene expression data from patients suffering from hematological malignancies. Mining available data from the MILE study using BloodSpot, we found a significant correlation between perturbed expression of ESCRT components and hematological disorders (Fig 1A). All 16 major classes of hematopoietic disorders were associated with misexpression of multiple core ESCRT components. Up or downregulation of a given ESCRT component was seen fairly uniformly across all lymphoid leukemias analysed, whereas it was sporadic in myeloid leukemias, suggesting that balanced hematopoiesis may require subtle control of various ESCRT components. Thus, lineage-specific roles of ESCRT components may be important for the maintenance of particular blood cell populations.

**Figure 1.**
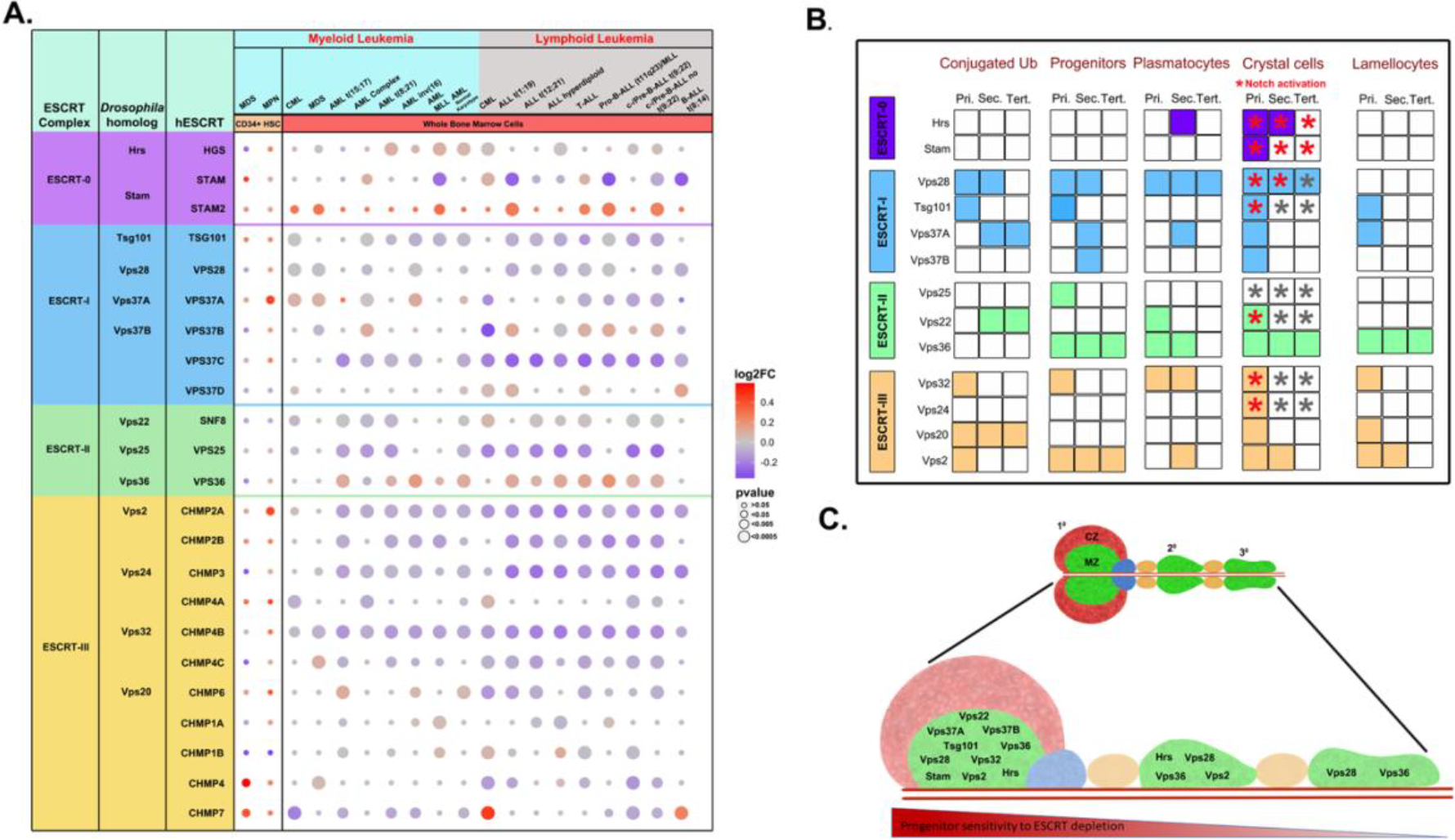
Dysregulation of ESCRT components perturbs hematopoietic homeostasis. **(A)** Change in expression levels of ESCRT components as found in CD34^+^ hematopoietic stem cells from MPN (Baumeister J et al., 2019) and MDS patients (Pellagatti A. et al., 2010) as well as in whole marrow cells obtained from leukemia patients (BloodSpot, MILE study) is visualized using a bubble plot, across all classes of leukemia described in the study. Shades of red indicate overexpression while shades of blue indicate downregulation of individual components. The intensity of color is correlative to the log2 fold change while the size of the individual bubble indicates the p-value. The left-most column lists the *Drosophila* homologs of the human ESCRT proteins. **(B)** Comprehensive summary chart of the effects of individual ESCRT depletion on the various aspects of hematopoiesis as indicated. Presence or absence of a phenotype is depicted by colored or white boxes, respectively. Red asterisk indicates Notch pathway activation and empty asterisk indicates no change, for components that were tested. **(C)** Schematic representation of lymph gland lobes from anterior to posterior (left to right). Green region (medullary zone) in primary and posterior lobes indicate the progenitor pool depleted of ESCRT components (13 genes from ESCRT-0, I, II and III). Red region (cortical zone) in the anterior lobe represents the mature hemocytes. ESCRT components mentioned in each lobe are those whose depletion affected crystal cell differentiation. Red monochrome heatmap indicates position-dependent progenitor sensitivity to depletion of ESCRT.

Data from the MILE study reflects the expression in a mixed population of bone marrow cells. Therefore, we also referred to individual microarray-based myelodysplatic syndrome (MDS) [21, 22] and myeloproliferative neoplasm (MPN) [20] studies done for CD34^+^ hematopoietic stem cells (HSCs) to understand whether ESCRT expression is perturbed in stem and progenitor cells. ESCRT-III components CHMP-5 and -7 were overexpressed in MDS whereas Vps37A of ESCRT-I and Vps2 of ESCRT-III in MPN. This suggests ESCRT-III may be more critical in the context of HSC maintenance and differentiation.

### ESCRT perturbation has non-uniform effects on ubiquitinated cargo sorting in the lymph gland progenitors

To test whether all ESCRT components have similar roles in cargo sorting in blood progenitors, we depleted each of the 13 core components individually in the lymph gland by RNAi mediated knockdown (KD) using the domeless (dome)-GAL4 driver that expresses in blood progenitors (*domeGal4*>*UAS ESCRT RNAi*). We checked expression and validated the knockdown of representative ESCRT components from each subunit and observed uniform expression across the lymph gland (Fig S1, see supplementary results). As ESCRT plays an active role in ubiquitinated cargo sorting in multiple tissue and cell types, the accumulation of ubiquitinated cargoes serves as a hallmark of dysfunctional ESCRT machinery and impaired endosomal protein sorting. Immunofluorescence analysis of conjugated ubiquitin (Ub) status (see methods) across all progenitor subsets (primary, secondary and tertiary lobes) showed a range of effects with the phenotype varying among ESCRT components within a given ESCRT complex and between complexes (Fig 1B, S2; see supplementary results).

Our analysis revealed non-uniform response of progenitors to perturbation of the cargo sorting machinery, depending on the ESCRT component that is depleted as well as the target progenitor population (Fig. 1B). Thus, we charted the spatiotemporal relation of Ub status to ESCRT depletion in LG progenitor subsets. We next tested whether this correlation reflects the response of progenitors to maintenance and differentiation cues and the developmental signaling pathways involved.

### ESCRT components play distinct roles in lymph gland progenitor maintenance

As ESCRT components regulate ubiquitinated cargo sorting in the blood progenitors, they might potentially regulate progenitor homeostasis. Hence, we checked whether depleting the 13 ESCRT components individually from dome^+^ progenitor subsets (marked by GFP expression) (*domeGal4>2XEGFP/+; UAS ESCRT-RNAi/+; +/+*) may affect progenitor status similar to the effect on Ub accumulation. As in the case of Ub status, a comprehensive analysis of progenitor fraction (see methods) in the primary, secondary, and tertiary lobes of the lymph gland showed a range of effects with the phenotype varying among ESCRT components within a complex and between complexes (Fig 1B, S3). Assessment of mitotically active nuclei and cell counts showed that ESCRT affects proliferation of blood progenitors (Fig. S4, S5, see supplementary results).

Since increased Ub accumulation indicates dysfunctional cargo sorting and this is likely a cause of progenitor loss, we compared Ub status on ESCRT KD with progenitor maintenance and found limited correlation. While knockdown of some ESCRT components [(Vps28, Tsg101 (ESCRT-I), Vps32 and Vps2 (ESCRT-III)] caused increased Ub and reduced progenitors in the primary lobe, others [(Vps25, Vps36 (ESCRT-II)] showed no change in Ub but progenitor numbers were reduced. Similarly, knockdown of Vps20 (ESCRT-III) had increased Ub but no effect on progenitors. Hence, in addition to Ub cargo sorting, ESCRT components Vps25 and Vps36 have a critical independent role in progenitor maintenance. Notably, there was no correlation at all between Ub increase and progenitor reduction in the youngest progenitors (tertiary lobe). This suggests that notwithstanding defects in cargo sorting, posterior progenitors remain less sensitive to perturbation, suggesting that they are maintained by other robust mechanisms (Fig 1C).

### ESCRTs downregulate Notch signaling to prevent crystal cell differentiation

As described earlier, 8 of the 13 core ESCRT components affect progenitor numbers and these effects are position-dependent. To examine the effect of progenitor status on hemocyte differentiation, we assessed plasmatocytes, crystal cells and lamellocytes in the lymph gland. Plasmatocytes, marked by P1 expression, make up about 95% of the differentiated hemocyte population but their numbers were increased only in a few cases upon ESCRT depletion (Fig 1B, S3; see supplementary results). Under steady state conditions each primary lobe harbors approximately 0-10 crystal cells, while they are generally absent from posterior lobes, even upon immune challenge [17]. Hence, we next analysed crystal cell status by checking expression of ProPO in the ESCRT depleted LGs.

Knockdown of 12 out of 13 ESCRTs increased crystal cell differentiation in the primary lobes (Fig 1B, S6), whereas Vps25 depletion had no effect. Secondary lobes were sensitive to depletion of Hrs, Vps28, Vps36 and Vps2 showing increased crystal cell numbers in all cases. Tertiary lobes showed increase in crystal cell numbers only on Vps28 or Vps36 depletion. This suggested that perturbation in the ESCRT machinery results in activation of signaling pathways that promote crystal cells. Notch pathway activation is a key requirement of crystal cell differentiation and Notch is a well-known target of ESCRT-mediated cargo sorting [6, 23-25]. Therefore, we examined the status of Notch pathway activation upon ESCRT depletion.

Increased crystal cell differentiation upon depletion of some ESCRT components suggests that they normally suppress Notch pathway activation. We focused our study on analysis of 8 ESCRT components [Hrs, Stam (ESCRT-0); Vps28, Tsg101 (ESCRT-I); Vps25, Vps22 (ESCRT-II); Vps32, Vps24 (ESCRT-III)] as these are well known for their role in NICD trafficking and Notch pathway activation [2, 7]. The Notch response element driving GFP (NRE-GFP) is a useful reporter to assess activation of Notch signaling. Upon KD in the lymph gland progenitors, all components tested caused upregulation of Notch signaling as interpreted by an increase in NRE-GFP positive cells in the primary lobe, except Vps25 (Fig 2A). Hrs, Stam and Vps28 depletion increased Notch activation in the posterior lobes (Fig 2A, S7A; see supplementary results). This is in concordance with the crystal cell differentiation phenotype as all of the ESCRT components except Vps25 cause increased crystal cell differentiation in the primary lobe upon knockdown. Our result also indicates that these 7 ESCRT components are indispensable for regulation of Notch signaling in the blood cell progenitors possibly with non-compensatory roles.

**Figure 2.**
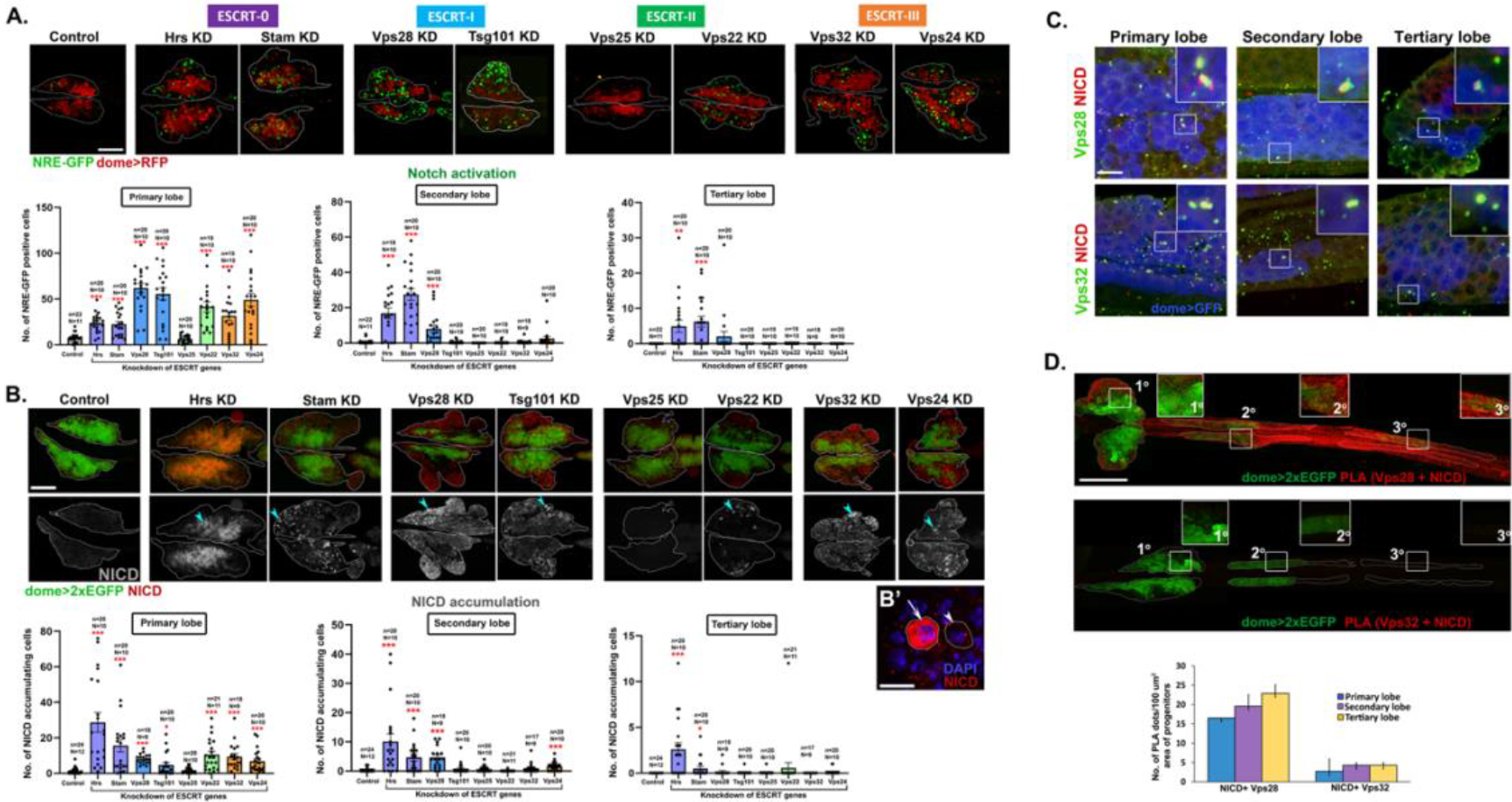
ESCRT components regulate Notch activation and NICD trafficking in the lymph gland. **(A)** Whole-mount larval lymph gland showing NRE-GFP+ve (Notch reporter) cells (green) and dome+ve progenitors (red) in primary lobes upon progenitor-specific knockdown of 8 representative ESCRT components indicated [Hrs, Stam (ESCRT-0); Vps28, Tsg101 (ESCRT-I); Vps25, Vps22 (ESCRT-II); Vps32, Vps24 (ESCRT-III)]. Scale bar: 100 µm. Bar diagrams show quantification of the number of NRE-GFP positive cells in primary, secondary and tertiary lobes upon knockdown of the same 8 ESCRT components. **(B)** Whole-mount larval lymph gland showing NICD expression (shown in red in the upper panel and in gray scale in the lower panel) in primary lobes upon progenitor-specific knockdown of the same 8 aforementioned ESCRT components. Progenitors are marked by dome>2xEGFP (green). Scale bar: 100 µm. **(B’)** Magnified view showing lymph gland hemocytes with (arrow) or without (arrowhead) NICD accumulation. Scale bar: 10 µm. **(C)** Immunostaining for NICD (red) and ESCRT components Vps28 and Vps32 (green) showing colocalization in dome+ve progenitors (blue) across primary, secondary and tertiary lobes of the lymph gland. Scale bar: 10 μm. **(D)** PLA dots (red) mark the interaction of NICD with Vps28 and Vps32 across three lobes of the lymph gland. Progenitors are marked by dome>2xEGFP. Insets show a magnified view of the progenitors from primary, secondary and tertiary lobes. Scale bar: 200 μm. Bar diagram shows quantification of the number of PLA dots per 100 μm2 area of the progenitors. N=5 larvae.

NICD transport and cleavage is required for the activation of Notch signaling. Accumulation of NICD may lead to aberrant activation of Notch signaling. Progenitor-specific knockdown of all tested ESCRT components, except Vps25, resulted in increase in the number of cells accumulating NICD in the primary lobe (Fig 2B). Also, this suggests a role for the majority of the ESCRT components in NICD trafficking, which may affect Notch signaling. However, the effects of perturbed Notch signaling on crystal cell differentiation vary in the posterior lobes, indicating position-dependent effects (see supplementary results). Since appropriate antibodies were available for Vps28 and Vps32, we checked for their expression and interaction with NICD. Immunolocalization and proximity ligation assay of ESCRT components (Vps28, Vps32) indicated uniform interaction of ESCRT with NICD (Fig 2C-D; see supplementary results). This suggests that additional mechanisms may effect progenitor maintenance, particularly in the posterior lobes. The absence of any phenotype due to Vps25 knockdown suggests compensatory mechanisms may regulate cargo trafficking and lineage-specific signaling activation, which is sufficient to maintain progenitors at steady state (Fig S8; see supplementary results).

### Position-dependent regulation of Notch signaling

3 out of the 8 ESCRT components tested [Hrs, Stam (ESCRT-0) and Vps28 (ESCRT-I)], upon knockdown caused increased Notch signaling activation in the secondary lobe (Fig 1B, 2A, S7A). Of these only Hrs and Vps28 knockdown resulted in increased crystal cell differentiation in the secondary lobe (Fig 1B, C, S6), suggesting additional mechanisms downstream of Notch activation prevent crystal cell differentiation in Stam KD. Interestingly, in the tertiary lobe, both Hrs or Stam knockdown resulted in Notch activation (Fig 2A, S7A) though neither resulted in crystal cell differentiation (Fig S6). In either case, tertiary lobe progenitors fail to differentiate, indicating their immature and refractile nature (Fig 1B, C, S6). Also, though Vps28 depletion does not significantly activate Notch signaling in the tertiary lobe it can promote crystal cell differentiation, suggesting a possible mechanism to downregulate Notch signaling likely after progenitors differentiate. However, we do see occasional increase in Notch activation in Vps28 KD tertiary lobes. Hence, crystal cell differentiation in the tertiary lobe could be under complex temporal regulation. Our analysis demonstrates the active role of ESCRT components in regulating Notch signaling, which may contribute to crystal cell differentiation and also differential response of the progenitor subsets.

### Notch regulation downstream of ESCRT is independent of Notch ubiquitination regulators

Knockdown of 4 components [Hrs, Stam (ESCRT-0), Vps28 (ESCRT-I) and Vps24 (ESCRT-III)] led to an increase in the number of NICD accumulating cells in the secondary lobe (Fig 2B, S7B). However, only Hrs and Stam knockdown resulted in NICD accumulation in the tertiary lobe. This is in concordance with the phenotype of Notch activation upon Hrs, Stam and Vps28 knockdown in the posterior lobes. Our results show that NICD accumulation and Notch pathway activation correlate perfectly with crystal cell differentiation upon ESCRT knockdown. However, the status of ubiquitinated cargo accumulation does not correlate strongly with crystal cell differentiation phenotype (Fig 3A). We hypothesized that Notch activation, triggered by ESCRT depletion in blood progenitors may be independent of the status of Notch ubiquitination. We analysed genetic interaction of ESCRT (Vps32) with Notch-specific ubiquitin ligase Deltex and deubiquitinase eIF3f1 in blood progenitors (Fig 3B-C, see supplementary results). Deltex or eIF3f1 depletion did not rescue increased Notch activation and crystal cell differentiation phenotype in Vps32 KD lymph gland. Hence, the role of ESCRT in Notch signaling may lie downstream of subtle post-translational regulatory mechanisms and could be independent of the canonical role of ESCRT in ubiquitinated cargo sorting.

**Figure 3.**
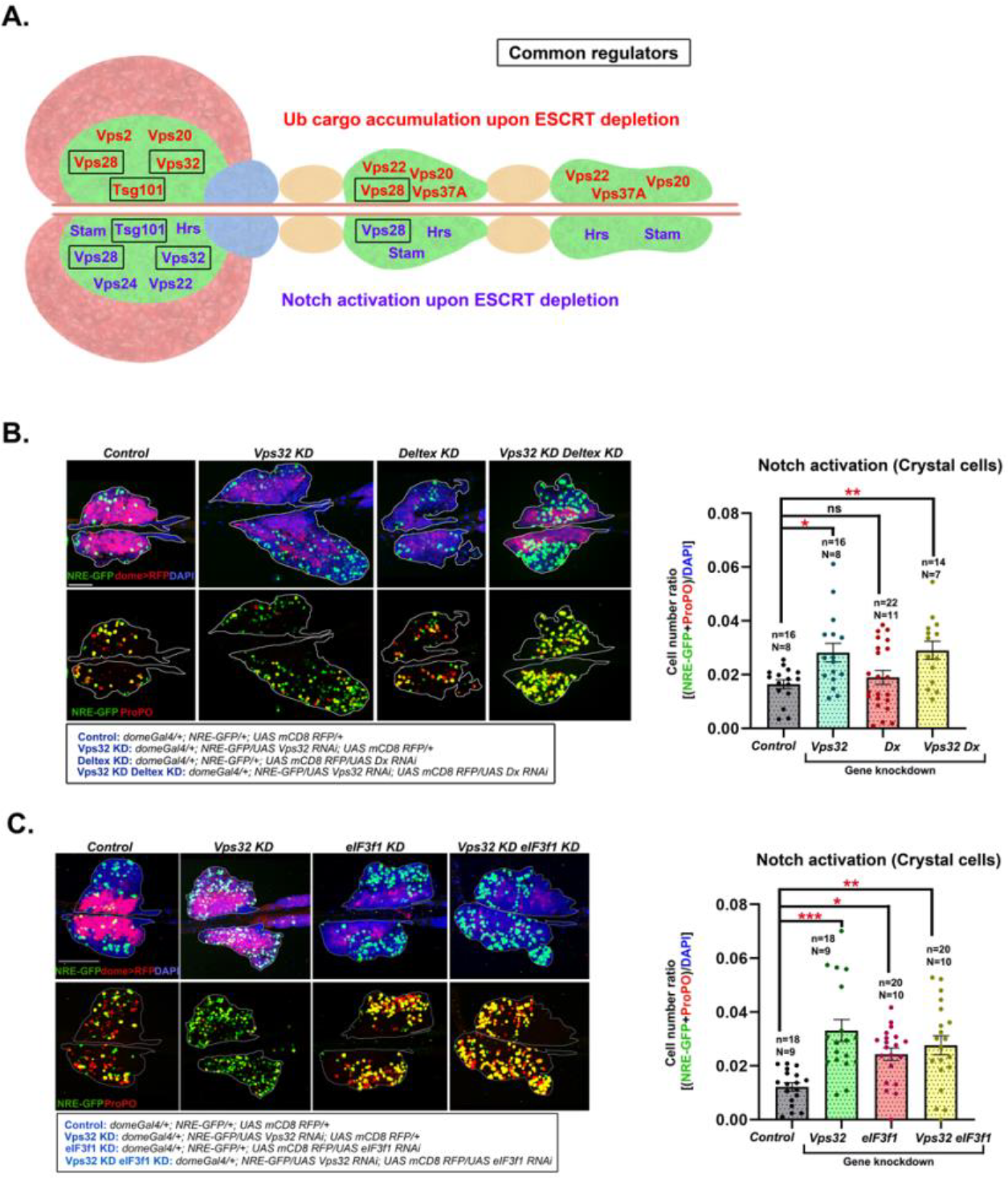
Non-canonical ESCRT functions may be at play for maintenance of hematopoietic homeostasis. **(A)** Schematic showing ESCRT components that affect ubiquitinated cargo sorting upon their depletion (in red) in the upper half and components that affect notch activation upon their depletion (in purple) in the lower half of the lymph gland schematic, in the respective lymph gland lobes. Black boxes indicate ESCRT proteins that affect both the processes in the respective lymph gland lobes. **(B)** Whole-mount larval lymph gland showing NRE-GFP and ProPO staining mark Notch activation and crystal cell differentiation, respectively in the primary lobe of control, Vps32 KD, eIF3f1 KD and Vps32 KD eIF3f1 KD lymph gland. Detailed genotypes are mentioned below the image panel of C and D. Bar diagrams show quantification of the number of NICD accumulating cells in primary, secondary and tertiary lobes. n indicates the number of individual lobes analysed and N indicates the number of larvae analysed. Error bars represent SEM. One-way ANOVA was performed to determine the statistical significance. *P<0.05, **P<0.01

### ESCRT depletion sensitizes progenitors for lamellocyte differentiation but does not confer survival advantage

Lamellocyte differentiation rarely occurs in the larva without any wasp infestation. However, progenitor-specific knockdown of 6 ESCRT components [Tsg101 and Vps37A (ESCRT-I); Vps36 (ESCRT-II); Vps32, Vps20 and Vps2 (ESCRT-III)] induced lamellocyte differentiation in the primary lobe, as visualized by Phalloidin (F-actin) staining, without any immune challenge (Fig 1B, S9). Knockdown of 2 components (Vps36 and Vps2) resulted in lamellocyte differentiation in the secondary lobe and only Vps36 knockdown triggered lamellocyte differentiation in the tertiary lobe. This indicates that the majority of the ESCRT components are not involved in suppressing lamellocyte differentiation in the refractile posterior progenitors at steady state. However, it is likely that KD progenitors may be more sensitive to immunogenic cues as compared to normal, unperturbed progenitors.

Wild type larvae are generally able to mount a sufficiently robust immune response that aids in their survival and eclosion against a low dose of wasp infestation. Systemic signals are generated upon wasp infestation and are received by the lymph gland progenitors [26], possibly through a complex extracellular matrix [17]. This results in lamellocyte differentiation in the primary lobe followed by disintegration and release of lamellocytes into circulation. Secondary and tertiary lobes are refractile to wasp infestation and do not form lamellocytes even upon immune challenge. Hence, we chose to test lamellocyte differentiation upon wasp infestation in-a] ESCRT KD that had no effect on lamellocyte formation (Vps25 KD) and b] ESCRT KD that caused lamellocyte differentiation only in the primary lobe (Vps32 KD). Knockdown of either Vps25 or Vps32 triggered lamellocyte differentiation across all progenitor subsets upon immune challenge with wasp (Fig 4A). This shows that Vps25 and Vps32 play essential roles in preventing all posterior progenitors from lamellocyte differentiation in response to a natural immune challenge. Further, loss of ESCRT sensitizes progenitors to systemic cues by unlocking differentiation programs.

**Figure 4.**
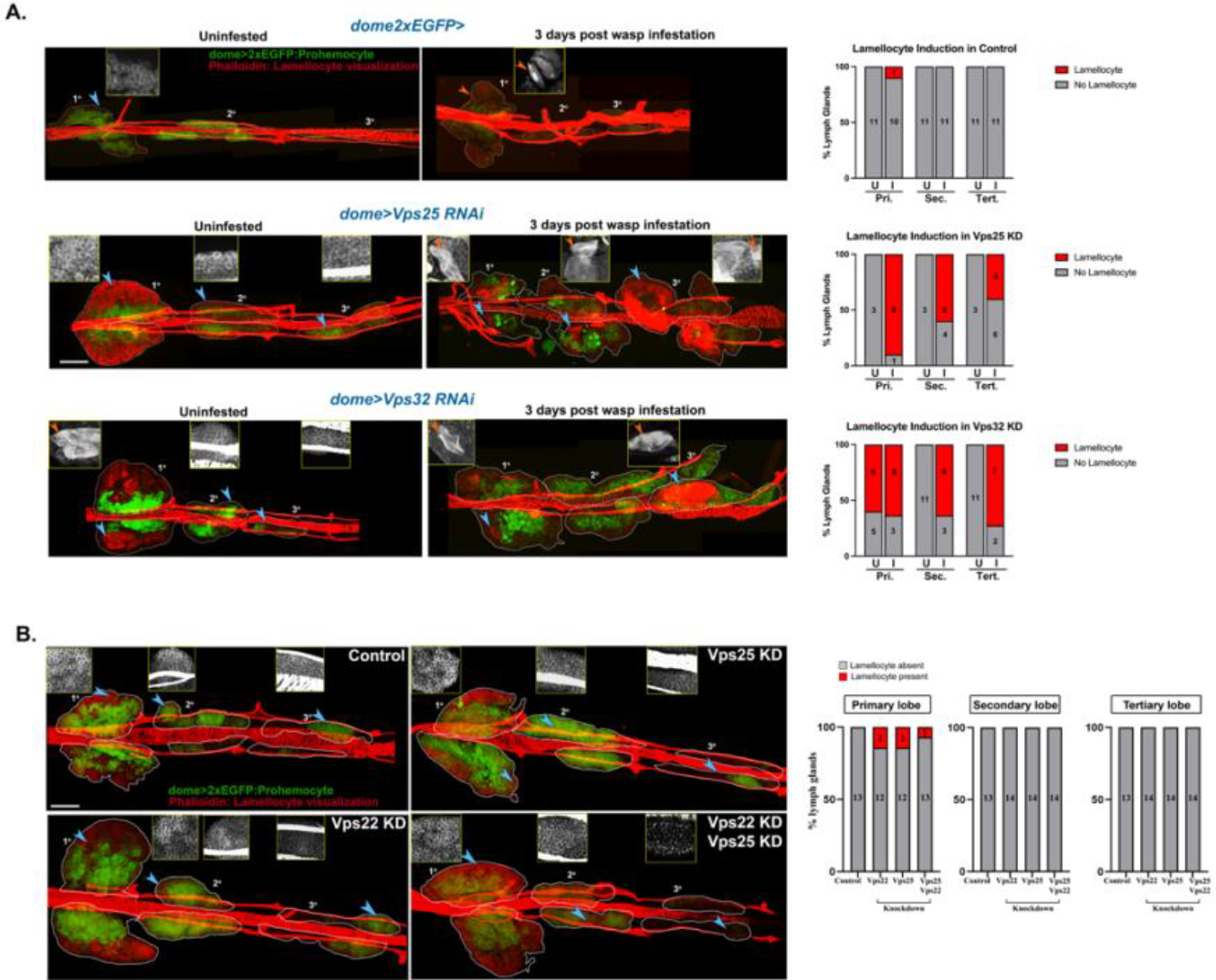
ESCRT components are non-compensatory in function and regulate prohemocyte sensitivity to immunological cues. **(A)** Whole-mount lymph glands of Vps25 KD (*domeGal4 UAS 2xEGFP; UAS Vps25 RNAi; +/+*) and Vps32 KD (*domeGal4 UAS 2xEGFP; UAS Vps32 RNAi; +/+*) larvae uninfested or 3 days after wasp infestation. Phalloidin staining shows presence of lamellocytes (marked by orange arrowhead in the inset). Bar diagram shows quantification of the percentage of lymph glands showing lamellocyte differentiation in primary, secondary and tertiary lobes upon knockdown of ESCRT components (Vps25 and Vps32), with and without immune challenge. Values in the columns indicate the number of larvae analysed for presence or absence of lamellocytes. **(B)** Whole-mount larval lymph gland showing Phalloidin staining (red) to visualise elongated morphology of lamellocytes upon progenitor-specific knockdown of ESCRT-II components [Vps22 (*domeGal4 UAS 2xEGFP; UAS Vps22 RNAi; +/+)*, Vps25 (*domeGal4 UAS 2xEGFP; UAS Vps25 RNAi; +/+)* and both Vps25 and Vps22 together (*domeGal4 UAS 2xEGFP; UAS Vps25 RNAi/UAS Vps22 RNAi; +/+*)]. Blue arrowheads mark the region from primary, secondary or tertiary lobes, magnified in the insets. The inset panel shows enlarged view of Phalloidin staining with or without lamellocytes. Scale bar: 100 µm. Bar diagram shows quantification of the percentage of lymph glands showing lamellocyte differentiation in primary, secondary and tertiary lobes upon knockdown of ESCRT-II components, without any immune challenge. Values in the columns indicate the number of larvae analysed for presence or absence of lamellocytes.

Our analyses showed that altered hematopoiesis due to blood progenitor-specific depletion of ESCRT does not considerably affect post-embryonic development until eclosion, indicating normal function of the differentiated hemocytes (Fig S10A, B). To test whether increased differentiation provides any survival advantage, we assessed survival in wasp challenged Vps25 KD, Vps32 KD and Vps36 KD larvae (domeMesoGal4 elavGal80-driven). ESCRT knockdown in the progenitor did not provide any survival advantage to the larvae as nearly all of the pupae failed to eclose with a lethal (moderate) dose of infection (Survival: Control 7.4%, Vps25 KD: 3%, Vps32 KD: 8.4%, Vps36 KD: 1.1%). Hence, *a priori* sensitization of progenitors does not promote survival upon immune challenge (Fig S10C). This could be due to rapid exhaustion of the progenitors in the late larval stage and insufficiency of lamellocyte count required to combat infection despite increased lamellocyte differentiation in the lymph gland. It is also likely that ESCRT depletion compromises lamellocyte ability to complete one or more steps involved in encapsulation, given the large number of signaling pathways that depend on ESCRT.

### Blood progenitor homeostasis does not require functional compensation between dispensable ESCRT components from the same subunit

Analysis of lamellocyte differentiation reflects functional diversity across ESCRT components (see Fig 1B, S9; supplementary results). Also, phenotypic diversity was observed within the same ESCRT subunit. Knockdown of ESCRT-II component Vps36 triggers spontaneous lamellocyte differentiation in steady state while depletion of the other two ESCRT-II components Vps25 or Vps22 does not affect lamellocyte differentiation. To confirm that Vps25 and Vps22 do not compensate each other in lamellocyte differentiation process, we generated larvae with double knockdown of Vps25 and Vps22 in the blood progenitor (*domeGal4 UAS2xEGFP/+; UAS Vps22 RNAi/UAS Vps25 RNAi;+/+*). Like the individual knockdown of Vps22 and Vps25, the double knockdown also did not trigger any significant lamellocyte differentiation under steady state (Fig 4B). This indicates that Vps22 and Vps25 do not take part in regulating lamellocyte differentiation. Hence, absence of steady state lamellocyte differentiation upon KD of certain ESCRT components is not due to functional compensation by other core components within the same ESCRT subunit. However, auxiliary components might functionally substitute core components in the blood progenitor which merit further investigation.

### ESCRT exhibits context dependent cell-autonomous and non-cell-autonomous roles in hematopoiesis

Progenitor-specific downregulation of ESCRT expression leads to accumulation of ubiquitinated cargoes. However, the majority of the ubiquitin accumulates in non-progenitor (Dome^-^) cells, suggesting a possible cell non-autonomous effect. The other possibility could be persistent ubiquitin accumulation even after the progenitors differentiate and lose Dome expression. We generated homozygous mutant mitotic recombinant clones for a representative ESCRT gene Vps32 (shrub), in progenitors. Vps32 is a terminally acting ESCRT (ESCRT-III) in the ubiquitinated cargo sorting pathway. Its depletion affects all blood cell lineages (Fig. 1B). Hence it serves as a good model to assess cell autonomous roles of endosomal protein sorting. Staining for conjugated ubiquitin revealed accumulation of ubiquitin aggregates in the homozygous mutant patch of the cells in the cortical zone (GFP^-^) indicating accumulation of cargo in blood progenitors depleted of ESCRT (Fig 5A). This suggests that despite a decrease in dome expression, ubiquitinated cargo sorting defects can persist in the ESCRT knockdown or knockout cells.

**Figure 5.**
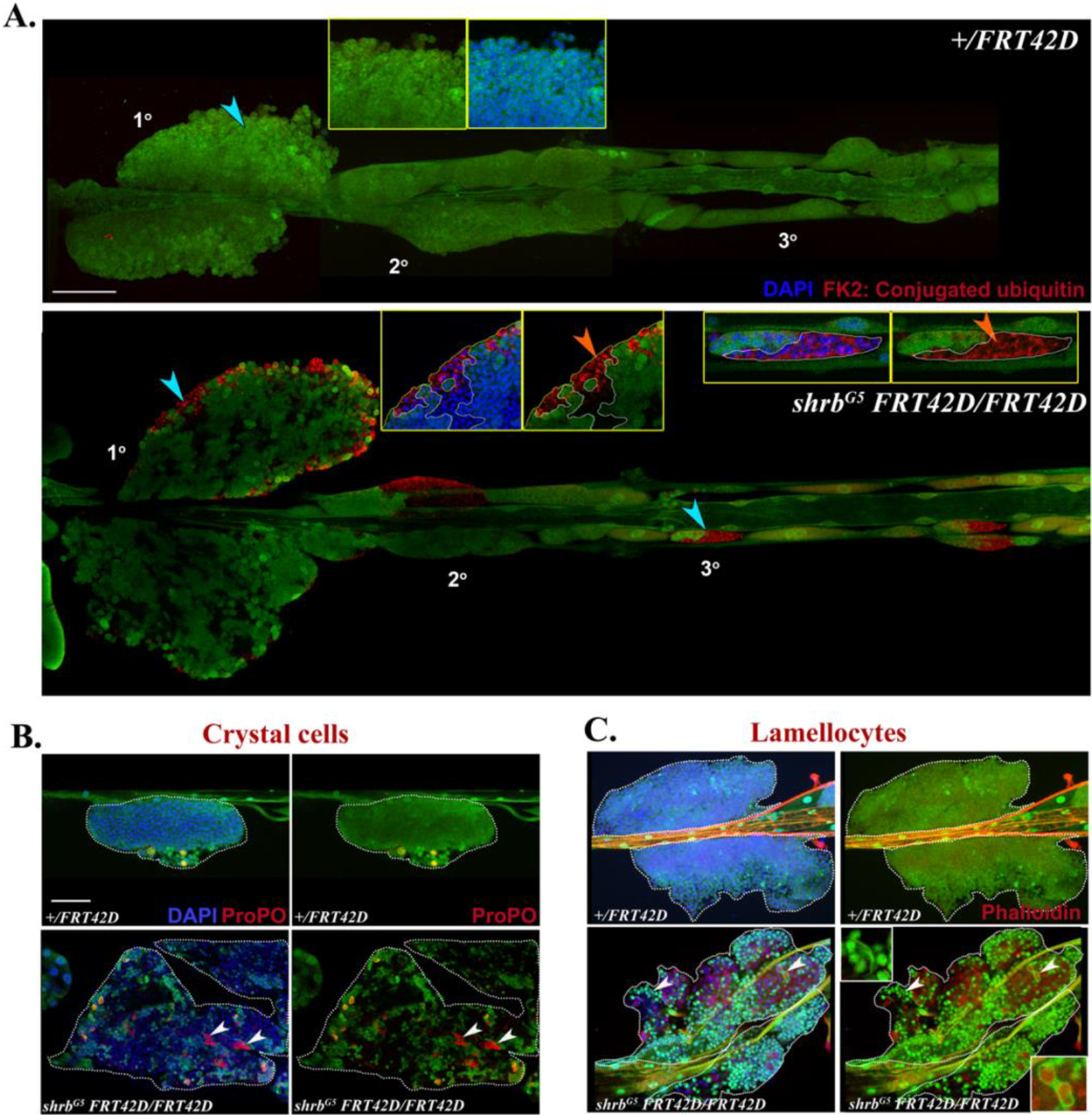
ESCRT cell-autonomously regulates ubiquitinated cargo sorting in the lymph gland progenitors but may regulate differentiation in non-autonomous manner as well. **(A)** Whole-mount lymph glands showing immunostaining for conjugated ubiquitin (red) in control (*domeGal4/+; neoFRT42D/+; UAS mCD8 RFP/+*) and progenitor-specific mutant clone of representative ESCRT component Vps32/shrub (*domeGal4/+; shrb*^*G5*^ *neoFRT42D/neoFRT42D; UAS mCD8 RFP/UAS FLP*). Area marked by blue arrowheads are magnified in the insets to show homozygous mitotic clones (GFP-ve patch, demarcated by dotted white line). Orange arrowheads indicate ubiquitin accumulation in the mutant cells. DAPI marks the nuclei. **(B)** Primary lobe of control and Vps32 mutant clone showing ProPO staining (red) to mark crystal cells. Arrowheads mark the crystal cells which are GFP-ve (homozygous mutant). **(C)** Phalloidin staining in the same genotypes shows GFP+ elongated and coalescing cells marked by arrowheads (insets) in the mutant clone. Scale bar in all image panels: 100 µm.

We analysed differentiation of blood cells in mitotic clones of ESCRT. ProPO staining showed both wild type and mutant origin of crystal cells as revealed by overlap with GFP expression in the mutant tissue (Fig 5B). As crystal cells are usually present in the lymph gland in low numbers, it is difficult to interpret the cell-autonomous origin of crystal cells from mutant progenitors. However, lamellocytes are completely absent in the control lymph gland at steady state (Fig S9). Phalloidin staining in the Vps32 mutant clone showed GFP-expressing elongated or coalescing cells, indicating the presence of lamellocytes and possibly their precursors (Fig. 5C). This suggests non-autonomous regulation of lamellocyte differentiation by ESCRT. Hence, ESCRT may regulate progenitor differentiation in both a cell-autonomous as well as cell non-autonomous manner. Since cell non-autonomous effects might hinge on cell-cell communication either through junctions or juxtacrine signalling, it suggests an overarching effect of ESCRT depletion to bring about differentiation of neighbouring cells to a particular lineage.

## Discussion

Cargo sorting by the ESCRT machinery is ubiquitously essential. Earlier work from our laboratory has shown that cargo sorting defects exhibited by NICD accumulation in Hrs+ endosomes, due to the knockout of a conserved regulator of hematopoiesis, Asrij, led to precocious differentiation and subsequent loss of blood progenitors. Mining available datasets of patient samples also indicated misexpression of ESCRT components in various hematological malignancies. Therefore, we asked whether ESCRTs play a decisive role in blood progenitor maintenance and lineage choice. Due to the pleiotropic subcellular roles of ESCRT proteins, it becomes challenging to assign specific cellular phenotypes to the ESCRT structures [27]. Our study now provides a means to test the complex functional diversity using a simple model where hematopoiesis can be sampled in its entirety, thereby allowing in-depth *in vivo* analysis.

We show that endosomal protein sorting actively maintains hematopoietic homeostasis. We uncover specific steps of endosomal protein sorting that potentially dictate blood progenitor maintenance. ESCRT-I remodels the endosomal membrane through budding and ESCRT-III carries out scission, to allow cargo sorting. Loss of ESCRT-I or ESCRT-III components result in progenitor differentiation suggesting that membrane budding and scission affects a wide range of signaling pathways across distinct progenitor subsets. Recent reports highlight the universal role of ESCRT-III, often in concert with ESCRT-I, in various membrane remodelling processes such as nuclear envelope reformation, lysosomal membrane repair, cytokinetic abscission, macroautophagy, exocytosis and lipid transport to mitochondria via lipid droplets [28, 29]. Whether such moonlighting functions of ESCRT-I and ESCRT-III impact signaling that determines progenitor maintenance merits further investigation.

Curiously, we observed drastic functional diversity of the highly conserved ESCRT-II components in progenitor differentiation. While Vps36 depletion affected all progenitor compartments across progenitor subsets, Vps25 depletion did not affect differentiation at steady state. Loss of Vps25 caused hyperproliferation in blood cells and failed to activate signaling pathways such as Notch, which are necessary for progenitor differentiation. Though Vps25 regulates endosomal protein sorting in epithelial tissues [25, 30], its redundancy in controlling differentiation suggests additional alternate routes for endosomal protein sorting in blood progenitors or a temporally regulated, developmental stage-specific role of Vps25 that has not yet been identified. Notably, though Vps25 is dispensable for steady state hematopoiesis, its depletion sensitizes all progenitors to differentiate upon immune challenge. This supports the possibility that the diverse roles of ESCRT may contribute to differential regulation of steady state and stress hematopoiesis.

Lymph gland progenitor subsets are functionally heterogeneous and show reduced sensitivity to differentiation cues from anterior to posterior [17, 18]. Posterior progenitors resist differentiation upon immune challenge suggesting that they have additional signal regulatory checkpoints. The mechanism by which progenitor subsets differentially respond to systemic cues remain largely unexplored. One possibility is that younger progenitors have inherently low levels of ubiquitination and protein turnover and hence show mild effects on ESCRT depletion. Depletion of Vps36 and Vps2 can trigger lamellocyte differentiation in refractile progenitors even without any immune challenge, indicating that they actively prevent differentiation. Also, while Hrs knockdown activates Notch signaling and crystal cell differentiation in posterior progenitors, Stam knockdown fails to trigger terminal differentiation to crystal cells despite Notch activation. This indicates existence of multiple checkpoints and highlights the complexity of mechanisms that progenitors may employ to maintain their identity. Elucidating expression and function at the single cell level may aid in an improved understanding of ESCRT-dependent lineage specification across these distinct progenitor subsets. However, positional information will be lost. Nevertheless, such candidates can be screened further for efficient modulation of vertebrate blood regeneration *in vitro* as well as *in vivo*.

Our study shows the distinct role of individual components of ESCRT in Notch signaling in blood progenitors. Expectedly, regulation of Notch signaling in lymph gland progenitors relies heavily on the ESCRT machinery. Our previous reports highlight the potential functional link of endosomal protein sorting with blood progenitor homeostasis [12, 13]. Though Hrs and Stam depletion promoted crystal cell differentiation in the lymph gland, it hardly affected plasmatocyte and lamellocyte differentiation. In epithelial tissue, though Hrs and Stam regulate Notch trafficking, they do not regulate Notch pathway activation and downstream phenotypes such as cell polarity and proliferation [7]. Similar mechanisms specific to hematopoietic tissue are not yet explored. It is notable that ALIX and its yeast homolog Bro1 can recognize non-ubiquitinated cargoes and sort them independent of ESCRT-0 [31]. Also, Bro1, ALIX and HD-PTP act as alternate bridging factors to ESCRT-II to mediate endosomal protein sorting in yeast and mammalian cells [32, 33]. Post-translational regulatory mechanisms may also render ESCRT components inactive [34].

The role of ESCRT in lamellocyte differentiation indicates the impact on upstream signaling pathways. EGFR signaling promotes plasmatocyte proliferation and lamellocyte differentiation. Also, downregulation of JAK/STAT and Hedgehog signaling leads to progenitor loss and lamellocyte differentiation [17, 26, 35]. While the role of ESCRT in EGFR and Hh signaling is known [2, 36, 37] the same in hematopoiesis is not dissected so far. Our analysis suggests that ESCRTs impact signaling pathways upstream of lamellocyte differentiation. Thus, we speculate that lineage-specific signaling activation could be achieved through modulation of specific ESCRT expression and function.

Deltex and eIF3f1 positively regulate Notch signaling in epithelial tissues through Notch ubiquitination [38-40]. However, downregulation of Dx/eIF3f1 failed to rescue Notch activation phenotype in ESCRT depleted progenitors. While other E3 ubiquitin ligases and deubiquitinases may possibly complement for Deltex or eIF3f1 depletion, signaling activation may not always depend on the status of cargo ubiquitination. For example, Vps36 depletion elicits a strong phenotype of differentiation without causing any change in ubiquitination. Further genetic interaction-based studies with other regulators of Notch signaling may reveal ESCRT-dependent mechanisms of Notch activation.

Loss of function mutation in ESCRT genes result in cell-autonomous cargo accumulation in *Drosophila* epithelial tissues [2, 6]. However, the known cell non-autonomous role of ESCRT in cell proliferation as well as neoplastic transformation, suggests altered intercellular communication and aberrant signaling activation in the neighboring cell population [2, 6, 25]. In concordance with the previous reports, we observed a cell-autonomous role of ESCRT in regulating ubiquitinated cargo sorting in the blood progenitors. However, analysis of lamellocyte differentiation suggests that ESCRT may affect lineage-specification non-autonomously. Both progenitor differentiation and proliferation influence blood cell homeostasis in the lymph gland. ESCRT depletion not only activates lineage-specific signaling pathways but also promotes blood cell proliferation. Cell type-specific increase in proliferation and enlargement of lymph gland lobes can affect the proportion of different hemocyte populations. Elucidating the interplay between ESCRT components and the mitogenic signaling machinery could reveal whether downregulation of mitotic potential may restore hematopoietic homeostasis.

Over 90% of hematological malignancies are associated with misexpression of one or more core ESCRT components. Hematological anomalies such as acute lymphoblastic leukemia (ALL), acute myeloid leukemia (AML), myelodysplastic syndromes, etc., stem from hyperproliferation and improper lineage choice caused by dysregulated activation of signaling pathways. While reports show the importance of ESCRT in the hematopoietic and immune system [15, 16] any functional analysis relating to progenitor maintenance and fate choice remained underexplored. The phenotypic diversity we observe highlights the role of the conserved ESCRT machinery in blood progenitor maintenance and differentiation. This may have implication in understanding blood disorders and designing target-specific therapeutic interventions.

## Materials and methods

### Fly stocks and genetics

*Drosophila melanogaster* stocks were maintained at 25°C as described previously [12]. The details of fly stocks, genetics and control genotypes used are in supplementary methods.

### Immunostaining analysis

*Drosophila* third instar larval lymph glands were dissected in PBS as described before and immunostained for microscopic analysis [19]. The detailed protocol and reagents used are in supplementary methods.

### MILE data collection and representation

For each ESCRT component, the expression values in each biological context (here, diseased and healthy bone marrow) were downloaded from BloodSpot (https://servers.binf.ku.dk/bloodspot/?gene=JAK2&dataset=all_mile). The expression values using all the probes for a particular gene were averaged and then the average expression values were normalized with the healthy bone marrow expression. Student’s t-test was performed for expression of each ESCRT component in each disease with healthy bone marrow using the BloodSpot plugin, and the p-values were noted. For the CD34^+^ hematopoietic stem cells (HSC) from Myeloproliferative Neoplasm (MPN) (GSE174060) and Myelodysplastic Syndrome (MDS) (GSE19429) patient samples, data was sourced from Baumeister J et al., 2019 and Pellagatti A. et al., 2010 respectively, using GEO [20, 21]. The datasets were analysed using GEO2R (LIMMA analysis with Benjamini-Hochberg FDR correction, adj. P-value <0.05). The log2 values of the obtained fold change were then plotted along with the p-value, in both cases, using the ggplot2 package in R. The resulting bubble plot indicates the log2FC values by the intensity of color, where red shows upregulation and blue downregulation, while the size of each bubble represents the p-value.

### Wasp parasitism assay

Wasp infestation was performed following standardised protocol as described in Rodrigues et al., 2021a [17]. Details are mentioned in supplementary methods.

### Quantification and statistical analysis

Blood cell differentiation was quantified as described in Ray et al., 2021 [18]. The details of quantification of various parameters and statistical analysis are mentioned in supplementary methods.

## Supporting information

Supplementary results

## Acknowledgements

This work was funded by grants to M.S.I from SERB and JC Bose fellowship, Department of Science and Technology, Government of India, LSRET grant from Department of Biotechnology and Jawaharlal Nehru Centre for Advanced Scientific Research (JNCASR), Bangalore. We thank JNCASR confocal facility for access and our laboratory members for helpful discussions.

## Author Contributions

A.R. and M.S.I. designed research; A.R. and Y.R. performed experiments; M.S.I. contributed reagents and materials; A.R., Y.R. and M.S.I. analysed the data and wrote the manuscript.

## References

1. Di Fiore, P.P. and M. von Zastrow, Endocytosis, signaling, and beyond. Cold Spring Harb Perspect Biol, 2014. 6(8).

2. Vaccari, T., et al., Comparative analysis of ESCRT-I, ESCRT-II and ESCRT-III function in Drosophila by efficient isolation of ESCRT mutants. Journal of Cell Science, 2009. 122(14): p. 2413–2423.

3. Alfred, V. and T. Vaccari, When membranes need an ESCRT: endosomal sorting and membrane remodelling in health and disease. Swiss Med Wkly, 2016. 146: p. w14347.

4. Radulovic, M., et al., ESCRT-mediated lysosome repair precedes lysophagy and promotes cell survival. EMBO J, 2018. 37(21).

5. Vietri, M., M. Radulovic, and H. Stenmark, The many functions of ESCRTs. Nat Rev Mol Cell Biol, 2019.

6. Herz, H.M., et al., Common and distinct genetic properties of ESCRT-II components in Drosophila. PLoS One, 2009. 4(1): p. e4165.

7. Tognon, E., et al., ESCRT-0 is not required for ectopic Notch activation and tumor suppression in Drosophila. PLoS One, 2014. 9(4): p. e93987.

8. Shravage, B.V., et al., Atg6 is required for multiple vesicle trafficking pathways and hematopoiesis in Drosophila. Development, 2013. 140(6): p. 1321–9.

9. Xia, P., et al., WASH is required for the differentiation commitment of hematopoietic stem cells in a c-Myc-dependent manner. J Exp Med, 2014. 211(10): p. 2119–34.

10. Reimels, T.A. and C.M. Pfleger, Drosophila Rabex-5 restricts Notch activity in hematopoietic cells and maintains hematopoietic homeostasis. J Cell Sci, 2015. 128(24): p. 4512–25.

11. Yu, S., F. Luo, and L.H. Jin, Rab5 and Rab11 maintain hematopoietic homeostasis by restricting multiple signaling pathways in Drosophila. Elife, 2021. 10.

12. Kulkarni, V., et al., Asrij maintains the stem cell niche and controls differentiation during Drosophila lymph gland hematopoiesis. PLoS One, 2011. 6(11): p. e27667.

13. Khadilkar, R.J., et al., ARF1-GTP regulates Asrij to provide endocytic control of Drosophila blood cell homeostasis. Proc Natl Acad Sci U S A, 2014. 111(13): p. 4898–903.

14. Avet-Rochex, A., et al., An in vivo RNA interference screen identifies gene networks controlling Drosophila melanogaster blood cell homeostasis. BMC Dev Biol, 2010. 10: p. 65.

15. Adoro, S., et al., Post-translational control of T cell development by the ESCRT protein CHMP5. Nat Immunol, 2017. 18(7): p. 780–790.

16. Liu, Y., et al., Membrane skeleton modulates erythroid proteome remodeling and organelle clearance. Blood, 2021. 137(3): p. 398–409.

17. Rodrigues, D., et al., Differential activation of JAK-STAT signaling reveals functional compartmentalization in Drosophila blood progenitors. Elife, 2021. 10.

18. Ray, A., K. Kamat, and M.S. Inamdar, A Conserved Role for Asrij/OCIAD1 in Progenitor Differentiation and Lineage Specification Through Functional Interaction With the Regulators of Mitochondrial Dynamics. Front Cell Dev Biol, 2021. 9: p. 643444.

19. Rodrigues, D., et al., Intact in situ Preparation of the Drosophila melanogaster Lymph Gland for a Comprehensive Analysis of Larval Hematopoiesis. Bio-protocol, 2021. 11(21): p. e4204.

20. Baumeister, J., et al., Early and late stage MPN patients show distinct gene expression profiles in CD34(+) cells. Ann Hematol, 2021. 100(12): p. 2943–2956.

21. Pellagatti, A., et al., Deregulated gene expression pathways in myelodysplastic syndrome hematopoietic stem cells. Leukemia, 2010. 24(4): p. 756–64.

22. Gorombei, P., et al., BCL-2 Inhibitor ABT-737 Effectively Targets Leukemia-Initiating Cells with Differential Regulation of Relevant Genes Leading to Extended Survival in a NRAS/BCL-2 Mouse Model of High Risk-Myelodysplastic Syndrome. Int J Mol Sci, 2021. 22(19).

23. Duvic, B., et al., Notch signaling controls lineage specification during Drosophila larval hematopoiesis. Curr Biol, 2002. 12(22): p. 1923–7.

24. Lebestky, T., S.H. Jung, and U. Banerjee, A Serrate-expressing signaling center controls Drosophila hematopoiesis. Genes Dev, 2003. 17(3): p. 348–53.

25. Vaccari, T. and D. Bilder, The Drosophila tumor suppressor vps25 prevents nonautonomous overproliferation by regulating notch trafficking. Dev Cell, 2005. 9(5): p. 687–98.

26. Morin-Poulard, I., A. Vincent, and M. Crozatier, The Drosophila JAK-STAT pathway in blood cell formation and immunity. JAKSTAT, 2013. 2(3): p. e25700.

27. Stempels, F.C., et al., Giant worm-shaped ESCRT scaffolds surround actin-independent integrin clusters. J Cell Biol, 2023. 222(7).

28. Olmos, Y., et al., ESCRT-III controls nuclear envelope reformation. Nature, 2015. 522(7555): p. 236–9.

29. Wang, J., et al., An ESCRT-dependent step in fatty acid transfer from lipid droplets to mitochondria through VPS13D-TSG101 interactions. Nat Commun, 2021. 12(1): p. 1252.

30. Herz, H.M., et al., vps25 mosaics display non-autonomous cell survival and overgrowth, and autonomous apoptosis. Development, 2006. 133(10): p. 1871–80.

31. Pashkova, N., et al., The yeast Alix homolog Bro1 functions as a ubiquitin receptor for protein sorting into multivesicular endosomes. Dev Cell, 2013. 25(5): p. 520–33.

32. Bissig, C. and J. Gruenberg, ALIX and the multivesicular endosome: ALIX in Wonderland. Trends Cell Biol, 2014. 24(1): p. 19–25.

33. Tabernero, L. and P. Woodman, Dissecting the role of His domain protein tyrosine phosphatase/PTPN23 and ESCRTs in sorting activated epidermal growth factor receptor to the multivesicular body. Biochem Soc Trans, 2018. 46(5): p. 1037–1046.

34. Tsunematsu, T., et al., Distinct functions of human MVB12A and MVB12B in the ESCRT-I dependent on their posttranslational modifications. Biochem Biophys Res Commun, 2010. 399(2): p. 232–7.

35. Mandal, L., et al., A Hedgehog- and Antennapedia-dependent niche maintains Drosophila haematopoietic precursors. Nature, 2007. 446(7133): p. 320–4.

36. Mattissek, C. and D. Teis, The role of the endosomal sorting complexes required for transport (ESCRT) in tumorigenesis. Mol Membr Biol, 2014. 31(4): p. 111–9.

37. Szymanska, E., N. Budick-Harmelin, and M. Miaczynska, Endosomal “sort” of signaling control: The role of ESCRT machinery in regulation of receptor-mediated signaling pathways. Semin Cell Dev Biol, 2018. 74: p. 11–20.

38. Matsuno, K., et al., Deltex acts as a positive regulator of Notch signaling through interactions with the Notch ankyrin repeats. Development, 1995. 121(8): p. 2633–44.

39. Moretti, J., et al., The translation initiation factor 3f (eIF3f) exhibits a deubiquitinase activity regulating Notch activation. PLoS Biol, 2010. 8(11): p. e1000545.

40. Hori, K., et al., Regulation of ligand-independent Notch signal through intracellular trafficking. Commun Integr Biol, 2012. 5(4): p. 374–6.

