## Supplementary results for "The Endosomal Sorting Complex, ESCRT, has diverse roles in blood progenitor maintenance, lineage choice and immune response"

### Supplemental Information

#### Supplementary Method:

#### *In situ* hybridisation

RNA *in situ* hybridisation was performed to check the expression of the ESCRT components against which antibodies are not available. The genes of interest (*Stam*, *Vps22*, *Vps25*, *Vps24*) were PCR amplified from genomic DNA using specific primer pairs as below, with the T7 promoter sequence incorporated in the reverse primer: *Stam* 5’ TTGTCACTGCCGATCTGTCC 3’ and 5’CCTGCTAATACGACTCACTATAGGGTTGACCCAGATAGCCACC 3’; *Vps22* 5’ACGTGATTTAGGTGACACTATAGTAGGCCTGGGAGCCATACAG 3’ and 5’ CCGTTAATACGACTCACTATAGGGTGCCAAAAATGCTCAATTTC 3’; *Vps25* 5’ CCCCAATTTAGGTGACACTATAGCGAAGAAACCAGACAGCAGC 3’ and 5’ TAATACGACTCACTATAGGGAAGAACTTAACGCCGTGGCTG 3’; *Vps24* 5’ GAGCCTGGTGCGCTATCC 3’ and 5’ GGCTTTAATACGACTCACTATAGGGTGCATCTCTTGCAGTTCCTC 3’

200ng-1µg of the amplicon was subjected to *in vitro* transcription using DIG-labelling mix (Roche, Switzerland) to generate DIG labelled RNA probes. Length of the probes are 644 bp (Stam), 503 bp (Vps25), 500 bp (Vps22) and 325 bp (Vps24). *In situ* hybridisation was performed as described in Benmimoun *et al*., 2015 [1].

#### RT qPCR

2-2.5 µg mRNA isolated from 100 lymph glands using Qiagen RNeasy kit was subjected to reverse transcription using Superscript (Invitrogen). 20ng cDNA was used for each qPCR reaction of *Stam*, *Vps25, Vps24* and *Rp49*. All SYBR green (Bio-Rad, USA)-based experiments were performed in triplicates. Relative fold change was estimated by normalising over Rp49. Primers used were- *Stam* 5’ ACTGAAAATGCGCCAAGTGC 3’ and 5’ CGGCAACAGTCTTGCTAGTC 3’; *Vps25* 5’ CCCTTCTTTACACTACAGCC 3’ and 5’ CTGGTCCCCAATGCTGAGAG 3’; *Vps24* 5’ AAGAGCAGGTGCAGGAGTGG 3’and 5’ CAAGAATGACGCAGGTGTCG 3’; *Rp49* 5’ CCGCTTCAAGGGACAGTATC 3’ and 5’ ACA ATC TCC TTG CGC TTC TTG 3’.

**Proximity ligation assay**

PLA was performed using the manufacturer's recommended protocol, as described before [2]. Rabbit plus and mouse minus probes were used along with orange DuoLink PLA kit (Merck, USA). Rabbit anti-Vps28 (Helmut Kramer, UT Southwestern Medical Center) or anti-Vps32 (Fen B. Gao, University of Massachusetts Medical School) and mouse anti-NICD (DSHB, USA) antibodies were used.

**Developmental analysis**

10 virgin females were crossed with 5 males in each vial containing cornmeal medium supplemented with yeast. After 8-10 hours of egg laying, the parents were flipped to a fresh vial for the next set. For each cross, 10 sets were made. The genotypes used were as follows:

With *DomeGal4 2xEGFP* driver- Control: *Dome-Gal4, UAS-2xEGFP>/Fm7a; +; +,* Vps25 KD: *Dome-Gal4, UAS-2xEGFP/+; UAS-Vps25 RNAi/+; +,* Vps36 KD: *Dome-Gal4, UAS-2xEGFP/+ ; UAS Vps36 RNAi/+; +,* Vps32 KD: *Dome-Gal4, UAS-2xEGFP/+; UAS-Vps32 RNAi/+; +.*

With Elav-Gal80;;Dome-mesoGFP> driver- Control: Elav-Gal80*; +;* Dome-mesoGFP>/+*,* Vps25 KD: Elav-Gal80*; UAS-Vps25 RNAi*/+*;* Dome-mesoGFP>/+*,* Vps36 KD: Elav-Gal80*; UAS Vps36 RNAi*/+*;* Dome-mesoGFP>/+*,* Vps32 KD: Elav-Gal80; *UAS-Vps32 RNAi/+;* Dome-mesoGFP>/+

Over 200 wandering third instar larvae were collected and sorted on the basis of GFP expression from each cross and age matched larvae were kept together. They were observed for pupariation and the subsequently eclosed flies were collected and their life span was recorded. The collected flies were flipped every alternate day to fresh vials, to avoid bacterial or fungal infections. Records were kept for larvae collected, percent pupariation, and death of adult flies on each day, and tabulated for analysis.

**Post-parasitism survival assay**

10 female L. boulardi wasps (Tina Mukherjee, InStem) from a running culture were introduced into vials with 40-50 late second instar larvae. Wasps were removed after 3 hours. Fly larvae were allowed to develop for 3 days till late third instar stage, and then GFP+ve larvae were collected for further observation. Number of eclosing adult flies was recorded for calculating percentage of survival.

The genotypes used were as follows:

With *DomeGal4 2xEGFP* driver- Control: *Dome-Gal4, UAS-2xEGFP>/Fm7a; +; +,* Vps25 KD: *Dome-Gal4, UAS-2xEGFP/+; UAS-Vps25 RNAi/+; +,* Vps36 KD: *Dome-Gal4, UAS-2xEGFP/+; UAS Vps36 RNAi/+; +.*

With Elav-Gal80;;Dome-mesoGFP> driver- Control: Elav-Gal80*; +;* Dome-mesoGFP>/+, Vps25 KD: Elav-Gal80*; UAS-Vps25 RNAi*/+*;* Dome-mesoGFP>/+, Vps36 KD: Elav-Gal80*; UAS Vps36 RNAi*/+*;* Dome-mesoGFP>/+, Vps32 KD: Elav-Gal80; *UAS-Vps32 RNAi/+;* Dome-mesoGFP>/+

### Supplementary Results

#### ESCRT expression is uniform across the lymph gland lobes.

Despite varying phenotype across progenitors upon ESCRT depletion, ESCRT components showed uniform expression across different lobes and developmental zones of the lymph gland, as assessed by immunofluorescence (IF) or RNA *in situ* hybridisation in wild type lymph gland (Fig S1A-C). This indicates that ESCRT function can be regulated post-transcriptionally to maintain spatiotemporally distinct progenitor subsets. *DomeGal4* driven knockdown of the ESCRT components was validated using immunofluorescence analysis, *in situ* hybridisation or RT-qPCR (Fig S1D-L).

#### Divergent requirement for ESCRT components in ubiquitinated cargo sorting in the *Drosophila* lymph gland progenitors.

Of the 13 core ESCRT components, 7 caused increased Ub in the LG when depleted. Interestingly, the effects were not uniform amongst progenitor subsets - 5 affected Ub status in the primary lobes, 4 in the secondary lobes and 3 in the tertiary lobes. Control LG showed low or no Ub in primary, secondary, and tertiary lobes (Fig 1B, S2). A similar trend was seen on depletion of ESCRT-0 components Hrs or Stam, with an occasional increase in Ub in primary lobes, which was not statistically significant (Figure S2A, B). In contrast, ESCRT-I, -II and -III depletion had effects on all lobes, though not all components affected the Ub status. Among ESCRT-I components (Vps28, Tsg101, Vps37A, Vps37B), depletion of Vps28 or Tsg101 very significantly increased Ub in the primary lobe, Vps28 and Vps37A affected the secondary lobe and Vps37A showed an increase in Ub in the tertiary lobe. Vps37B depletion had a mild non-significant effect on the Ub status of the LG. ESCRT-II components Vps25, Vps22 and Vps36 had no effect on the primary lobe.

However, Vps22 depletion caused a dramatic increase in Ub in the secondary and tertiary lobes, where Vps25 and Vps36 depletion had no effect. Finally, depletion of ESCRT-III components (Vps32, Vps20 and Vps2) caused a significant increase in Ub in the primary lobes whereas secondary and tertiary lobes were sensitive only to Vps20 depletion.

**ESCRT components play distinct roles in lymph gland progenitor maintenance.**

In controls, anterior lymph gland lobe progenitors are restricted to the inner medullary zone (MZ) while the posterior lobes are composed almost entirely of progenitors [6]. ESCRT-0 (Hrs, Stam) knockdown did not show any significant change in progenitor status indicating non-essential roles for these in progenitor maintenance (Fig S3A, B). Depletion of ESCRT-I components Vps28 and Tsg101 caused reduction in progenitor fraction in the primary lobes whereas secondary lobe progenitors were reduced by depletion of Vps28, Vps37A or Vps37B but not of Tsg101. Interestingly, ESCRT-I components had no effect on tertiary lobe progenitors. Among ESCRT-II components, Vps25 had an effect on proliferation causing an absolute increase in primary lobe cell numbers with a concomitant decrease in progenitor fraction (Fig S3A, B; S4A, C). Vps22 also did not affect LG progenitor fraction. In contrast, Vps36 drastically reduced progenitor fraction in all LG lobes, with phenotype severity increasing from anterior to posterior. ESCRT-III had very restricted effects on progenitors with Vps32 KD causing a reduction only in anterior progenitors, Vps2 KD reduced both anterior and posterior progenitors and Vps24 and Vps20 had no effect (Fig. S3A, B).

**Older progenitors are more prone to plasmatocyte differentiation upon ESCRT depletion**

Plasmatocytes, marked by P1 expression, make up about 95% of the differentiated hemocyte population. In the LG, they are restricted to the cortical zone of the primary lobe, with occasional P1 positive cells seen in posterior lobes. Reduced progenitor numbers are expected to be accompanied by an increase in the plasmatocyte population due to differentiation. Enumeration of P1 positive cells in the ESCRT KD LG (*domeGal4 UAS 2XEGFP> UAS ESCRT RNAi*) showed an expected increase in the plasmatocyte fraction of the primary lobe for Vps28, Tsg101, Vps36 and Vps32, where the progenitor fraction was mostly reduced (Fig 1B, Fig S3A, C). However, Vps25 depletion had no apparent effect on differentiation. This could be due to the failure of progenitors to terminally differentiate into plasmatocytes or due to non-autonomous over-proliferation of the intermediate population. Additionally, Vps22 KD also showed increased plasmatocyte numbers though there was no significant effect on the progenitor fraction, suggesting possible non-autonomous over-proliferation or exhaustion of the intermediate population. The remaining ESCRT components had no effect on primary lobe plasmatocytes.

Interestingly, KD of ESCRT-0 component Hrs and ESCRT-III component Vps32, that had no effect on progenitors, caused an increase in plasmatocyte numbers only in the secondary lobes. This indicates that though there is no effect as assessed by progenitor marker analysis, Hrs or Vps32 depletion has sensitized the tissue to respond to proliferation and differentiation cues. Along similar lines, Vps36 and Vps2 depletion drastically reduced the secondary progenitor fraction and increased plasmatocyte differentiation. Except for Vps28, ESCRT KD did not induce plasmatocytes in the tertiary lobes, even when progenitors were lost (e.g. Vps36 KD and Vps2 KD).

**ESCRT components play distinct roles in regulating mitotic potential across different blood progenitor subsets.**

Altered cell proliferation can contribute to perturbed tissue homeostasis. Our analysis of blood cell differentiation is based on the estimation of the cell fraction, which could be an outcome of not only progenitor differentiation but also proliferation of individual blood cell types. ESCRT genes act as tumor suppressors in epithelial tissues by inhibiting Notch-dependent hyperplastic and neoplastic overgrowth [3-5]. We tested whether downregulation of ESCRT expression can impact proliferation of blood cells. Analysis of 8 selected ESCRT components showed that progenitor-specific knockdown of 3 ESCRT components [Vps28 (ESCRT-I), Vps22 (ESCRT-II) and Vps32 (ESCRT-III)] led to an increase in the number of nuclei with high mitotic potential in the primary lobe as revealed by high level of phosphorylated Histone H3 (Fig S4A, B). Vps32 knockdown also caused an increase in the size of the primary lobe as interpreted by the number of nuclei (Fig S4A, C). However, Vps28 and Vps22 knockdown did not affect the overall size of the primary lobe. This suggests that cells may not have actively divided in the Vps28 and Vps22 depleted primary lobes in spite of increase in the mitotic potential. Though Tsg101 and Vps25 knockdown did not increase the number of H3P high nuclei in the primary lobe, the overall size of the primary lobe increased. This suggests possible early developmental stage-specific effects on cell proliferation due to ESCRT depletion.

Depletion of Vps22 (ESCRT-II) resulted in increase in the mitotic potential in the secondary lobes (Fig S4A, B). Depletion of 6 components [Hrs, Stam (ESCRT-0); Vps28, Tsg101 (ESCRT-I); Vps22 (ESCRT-II) and Vps32 (ESCRT-III)] however resulted in increase in cell number in the secondary lobes (Fig S4A, C). This again reflects temporal regulation of mitotic potential and cell proliferation upon knockdown of various ESCRT components. On the other hand, Vps25 (ESCRT-II) and Vps24 (ESCRT-III) knockdown reduced mitotic potential in the secondary lobe though the overall size of the secondary lobe remained unaffected (Fig S4A-C). This indicates positive regulation of mitotic potential by Vps25 and Vps24 in the secondary lobe.

None of the ESCRT KD showed increased mitotic potential in the tertiary lobe. Rather, proliferative potential decreased in the tertiary lobe upon knockdown of 4 components [Vps28, Tsg101 (ESCRT-I); Vps25 (ESCRT-II) and Vps24 (ESCRT-III)] (Fig S4A, B). However, tertiary lobe size increased upon knockdown of 6 components [Hrs, (ESCRT-0); Vps28 and Tsg101 (ESCRT-I); Vps22 (ESCRT-II); Vps32 and Vps24 (ESCRT-III)] (Fig S4A, C) suggesting hyperplasia. In summary, depletion of ESCRT components promote proliferation but in a developmentally regulated manner.

Increased differentiation or proliferation can cause disintegration of the primary lobe or appearance of tumorous bulges resembling neoplastic overgrowth. To assess the change in morphology of the lymph gland lobes upon depletion of ESCRT components, we categorized lymph glands into three groups based on the appearance of lobe margin: regular, irregular/disintegrated, and tumorous bulge. While the majority of the control lymph gland primary lobes showed a regular boundary, knockdown of 6 ESCRT components [Hrs (ESCRT-0); Vps28, Tsg101 (ESCRT-I); Vps25, Vps22 (ESCRT-II) and Vps32 (ESCRT-III)] resulted in tumorous outgrowth in the primary lobe (Fig S5 A, B). Also, Hrs and Vps28 knockdown resulted in a significant increase in primary lobe disintegration as revealed by the irregular boundary. Vps28 and Vps32 knockdown resulted in significant increase in tumorous overgrowth in the secondary and tertiary lobe. Our analyses show that altered mitotic potential and cellular proliferation due to ESCRT depletion can contribute to altered blood cell homeostasis.

**ESCRT components uniformly interact with NICD across progenitor subsets.**

As progenitor subsets respond differentially upon ESCRT depletion, we investigated whether co-expressing ESCRT components may differentially interact with endosomal cargoes in these different progenitor subpopulations. We chose two representative components, Vps28 and Vps32 that differentially regulate cargo sorting and progenitor homeostasis in the lymph gland. While Vps28 knockdown results in cargo accumulation and signaling activation in anterior as well as posterior subsets of progenitors, Vps32 depletion primarily affects anterior progenitor homeostasis. Immunostaining-based analysis showed co-localization of Vps28 and Vps32 with NICD across all progenitor subsets (Fig 2C). Using

Proximity ligation assay (PLA) to assess physical interaction, we found that while NICD interacted with Vps28 in progenitors of all three lobes as expected, direct physical interaction was negligible between Vps32 and NICD and did not vary across lobes (Fig 2D). Our results suggest that the phenotypic diversity across lobes may not arise due to differential expression or subcellular localization of individual ESCRT. Additional regulators may cause differential sorting efficiency by ESCRT components across progenitor subsets and merit further exploration.

**ESCRT regulates Notch activation in blood progenitors independent of Deltex and eIF3f1.**

Ubiquitinated cargo accumulation such as Notch intracellular domain (NICD) accumulation is a hallmark of ESCRT dysfunction. Yet, crystal cell differentiation phenotypes seen on ESCRT KD overlap only partially with increased ubiquitination (Fig 3A). Several ubiquitin ligases and deubiquitinases regulate Notch activation in various contexts [7]. Ubiquitin ligases such as Mindbomb, Neuralized, d-cbl, and Archipelago ubiquitinate the Notch-specific ligand Delta [8-10]. The Notch-specific E3 ubiquitin ligase Deltex positively regulates Notch pathway in a ligand-independent manner [11]. Also, Deltex acts in synergy with ESCRT-III component Vps32 in the *Drosophila* wing disc to fine-tune Notch signaling [12]. The Notch-specific deubiquitinase eIF3f1 acts downstream of Deltex and promotes γ-secretase-dependent cleavage of NICD from the endosome surface, thus upregulating Notch signaling [7]. As both Deltex and eIF3f1 positively regulate Notch signaling in *Drosophila* tissues by directly regulating the ubiquitination or deubiquitination of Notch [7, 12, 13], we probed into any possible genetic interaction between ESCRT and Deltex or eIF31 in blood progenitors, which may regulate Notch signaling.

Vps32 knockdown in the progenitors upregulates Notch signaling and crystal cell differentiation. However, progenitor-specific knockdown of Deltex (*domeGal4/+; UAS Vps32 RNAi/NRE-GFP; UAS Dx RNAi/UAS mCD8 RFP*) (Fig 3B) or eIF3f1 (*domeGal4/+; UAS Vps32 RNAi/NRE-GFP; UAS eIF3f1 RNAi/UAS mCD8 RFP*) (Fig 3C) failed to rescue the phenotype of Vps32 knockdown (*domeGal4/+; UAS Vps32 RNAi/NRE-GFP; +/UAS mCD8 RFP)*. Deltex or eIF3f1 knockdown maintained high Notch activation in Vps32 knockdown lymph glands. Our result indicates that Notch regulation downstream of ESCRT is independent of Notch ubiquitination regulators, Deltex or eIF3f1, at least for the ESCRTs tested. This might explain why depletion of some ESCRT components, despite showing no significant accumulation of cargo in ubiquitinated state can promote Notch signaling and crystal cell differentiation.

**Vps25 is dispensable for progenitor maintenance**

Vps25 knockdown did not affect the status of ubiquitination, progenitor maintenance or differentiation to any particular blood cell lineage despite its expression in the lymph gland. To further verify this, we generated lymph gland progenitor-specific homozygous clones of Vps25 loss of function mutation ($Vps25^{A3}$). There was no accumulation of ubiquitin aggregates or any change in the status of the progenitor (Fig S8A), plasmatocyte (Fig S8B), crystal cell (Fig S8C) and lamellocyte differentiation (Fig S8D). However, the mutant lymph glands showed enlargement of the primary lobe, suggesting possible increase in blood cell proliferation upon loss of Vps25. Also, phalloidin staining revealed appearance of binucleate, large cells and also very small cells occasionally, along with increase in F-actin content in some patches of the tissue, mostly in a cell autonomous manner (visible in GFP negative area of the tissue). Hence, Vps25 possibly inhibits uncontrolled cell proliferation and may contribute to critical steps of cell division that may dictate cell shape, number and polarity.

**ESCRT depletion causes spontaneous lamellocyte differentiation across progenitor subsets.**

Lamellocyte differentiation rarely occurs in the larva without any wasp infestation. However, progenitor-specific knockdown of 6 ESCRT components [Tsg101 and Vps37A (ESCRT-I); Vps36 (ESCRT-II); Vps32, Vps20 and Vps2 (ESCRT-III)] induced lamellocyte differentiation in the primary lobe, as visualized by Phalloidin (F-actin) staining, without any immune challenge (Fig 1B, S9A, B). Knockdown of 2 components (Vps36 and Vps2) resulted in lamellocyte differentiation in the secondary lobe and only Vps36 knockdown triggered lamellocyte differentiation in the tertiary lobe (Fig 1B, S9A, B ). This indicates that the majority of the ESCRT components are not involved in suppressing lamellocyte differentiation in the refractile posterior progenitors at steady state. However, it is likely that KD progenitors may be more sensitive to immunogenic cues as compared to normal, unperturbed progenitors.

**ESCRT depletion in hematopoietic compartments does not affect development and lifespan**

We performed a developmental analysis in ESCRT knockdown flies with or without wasp challenge. Hemocytes are essential for normal pupariation and eclosion in *Drosophila* (Stephenson et al., 2022). Vps25 knockdown reduced pupariation (Control: 98.41 %, Vps25 KD: 93.73 %) and eclosion (Control: 91.64

%, Vps25 KD: 88.13 %) as well as survival after eclosion. To abrogate the possibility of neuron-specific effect of Vps25 depletion on development and survival, we additionally used domeMesoGal4;elavGal80 driver to prevent Vps25 knockdown in the brain. This led to an increase in pupariation but showed only a mild effect on eclosion and led to reduced lifespan of adult flies, similar to domeGal4>Vps25 KD (Pupariation- control: 73.33%, Vps25 KD: 93.68%; eclosion- control: 92.3%, Vps25: 95.28%) (Fig S10B). Vps32 knockdown using domeGal4 showed normal pupariation, however, leading to death in the pupal stage (Fig S10A). Suppressing knockdown in neurons using domeMesoGal4 elavGal80 showed no effect on pupariation or eclosion and caused a mild reduction in adult fly survival between 30-60 days (Fig S10B). domeGal4-driven knockdown of Vps36 leads to spontaneous differentiation to all lineages across progenitor subsets. However, it did not affect the rate of pupariation (Control: 95.79%, vps36 KD: 92.54%) or eclosion (Control: 94.61%, Vps36 KD: 94.44%) (Fig S10A). Our analyses show that altered hematopoiesis due to blood progenitor-specific depletion of ESCRT does not considerably affect post-embryonic development until eclosion, indicating normal function of the differentiated hemocytes.

**Supplementary References:**


1. Benmimoun, B., Polesello, C., Haenlin, M., and Waltzer, L. (2015). The EBF transcription factor Collier directly promotes Drosophila blood cell progenitor maintenance independently of the niche. Proc Natl Acad Sci U S A *112*, 9052-9057.
2. Khadilkar, R.J., Rodrigues, D., Mote, R.D., Sinha, A.R., Kulkarni, V., Magadi, S.S., and Inamdar, M.S. (2014). ARF1-GTP regulates Asrij to provide endocytic control of Drosophila blood cell homeostasis. Proc Natl Acad Sci U S A *111*, 4898-4903.
3. Hariharan, I.K., and Bilder, D. (2006). Regulation of imaginal disc growth by tumor-suppressor genes in Drosophila. Annu Rev Genet *40*, 335-361.
4. Vaccari, T., and Bilder, D. (2005). The Drosophila tumor suppressor vps25 prevents nonautonomous overproliferation by regulating notch trafficking. Dev Cell *9*, 687-698.
5. Horner, D.S., Pasini, M.E., Beltrame, M., Mastrodonato, V., Morelli, E., and Vaccari, T. (2018). ESCRT genes and regulation of developmental signaling. Semin Cell Dev Biol *74*, 29-39.
6. Rodrigues, D., Renaud, Y., VijayRaghavan, K., Waltzer, L., and Inamdar, M.S. (2021). Differential activation of JAK-STAT signaling reveals functional compartmentalization in Drosophila blood progenitors. Elife *10*.

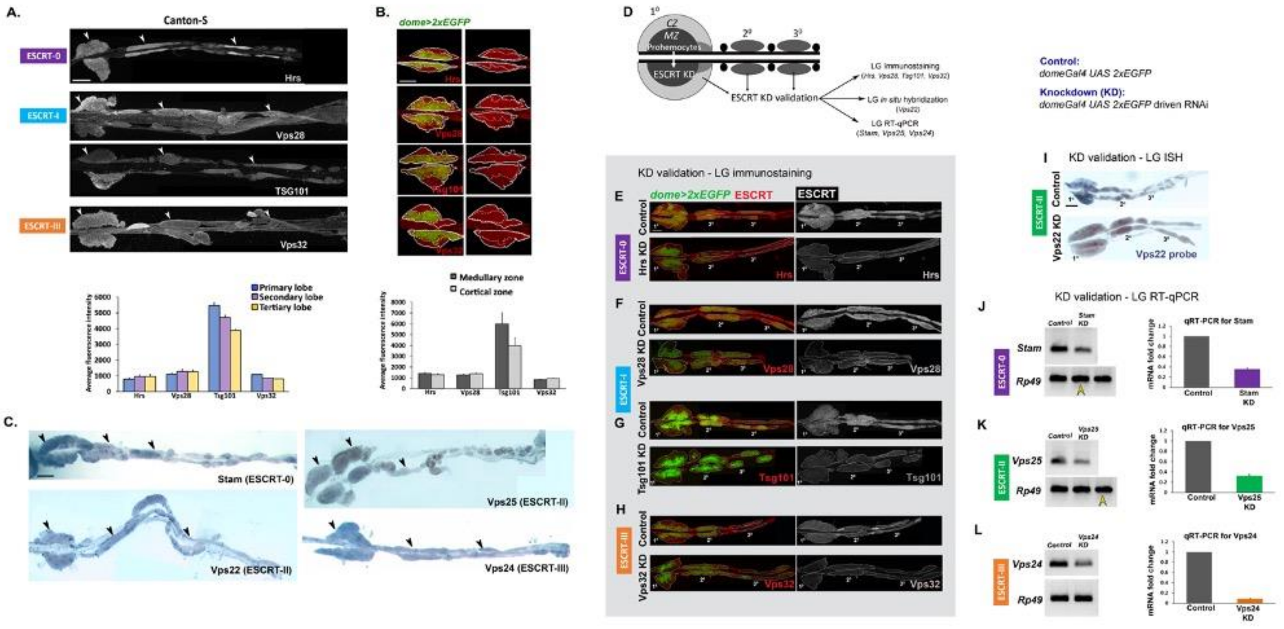


**Supplementary Figure S1**

**Supplementary Figure S1. ESCRT is uniformly expressed in the lymph gland and downregulated after RNAi.**

**(A)** Immunofluorescence microscopy of whole mount Canton-S lymph glands showing expression of ESCRT components Hrs (ESCRT-0), Vps28 and Tsg101 (ESCRT-I) and Vps32 (ESCRT-III) across different lobes of the lymph gland. Arrowheads mark the primary, secondary and tertiary lobes. Bar diagram shows quantification of mean fluorescence intensity for immunostaining of ESCRT components across three lobes. **(B)** dome>2xEGFP+ve region marks prohemocytes in the medullary zone demarcated by dotted line in the primary lobe. Immunostaining is shown for Hrs, Vps28, Tsg101 and Vps32. Bar diagrams show quantification and comparison of mean fluorescence intensity of a given component in dome>2xEGFP+ve medullary zone and dome>2xEGFP-ve cortical zone. **(C)** RNA *in situ* hybridisation shows expression of ESCRT components Stam (ESCRT-0), Vps22 and Vps25 (ESCRT-II) and Vps24 (ESCRT-III) at transcript level across different lobes. Arrowheads mark the different lobes. Scale bar: 100 µm. N>5 larvae with each individual lobes analysed. Error bars in the graph represent SEM. One-way ANOVA was performed to determine the statistical significance. **(D-H)** Validation of *domeGal4*-driven knockdown of ESCRT components was performed using immunofluorescence microscopy*, in situ* hybridisation and RT-qPCR (D). Immunostaining using respective antibodies shows knockdown of ESCRT component Hrs (ESCRT-0) (E), Vps28 (F) and Tsg101 (ESCRT-I) (G) and Vps32 (ESCRT-III) (H). **(I)** RNA *in situ* hybridisation shows knockdown of Vps22 (ESCRT-II). **(J-L)** RT-qPCR from lymph gland validates knockdown of Stam (ESCRT-0), Vps25 (ESCRT-II) and Vps24 (ESCRT-III). Scale bar: 100 µm. N>5 larvae for IF and N=10 larvae for ISH-based validation. RT-qPCR was performed in triplicates using RNA isolated from 100 lymph glands for each genotype. Error bars represent SEM.

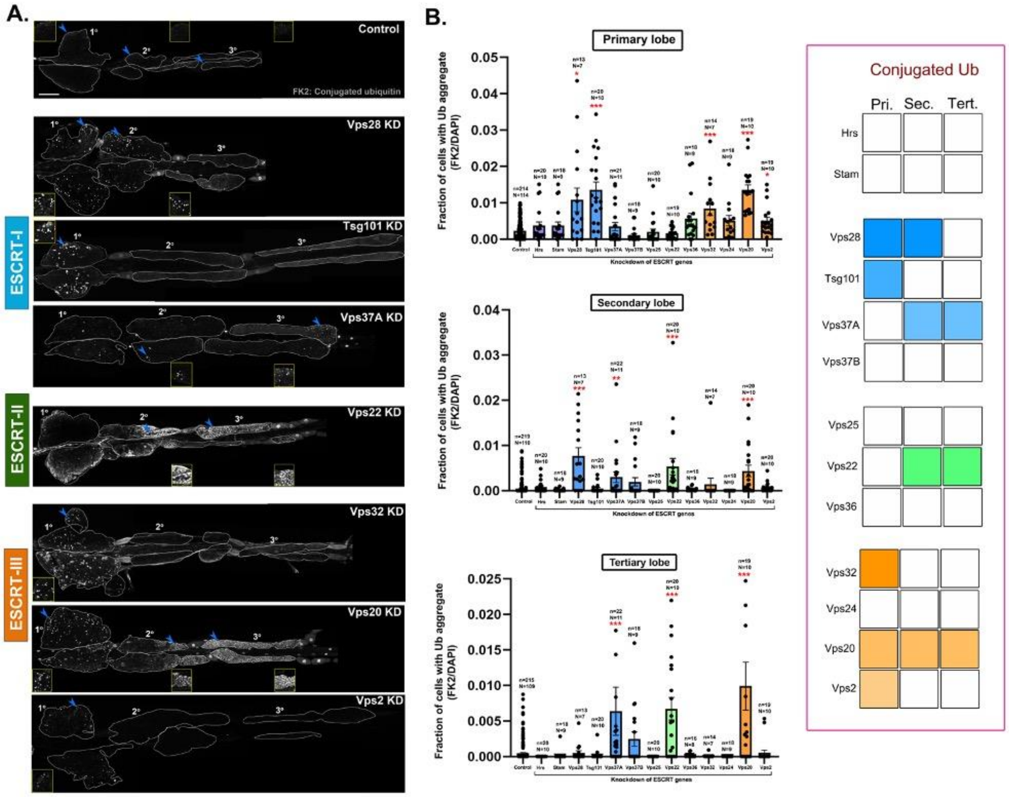


**Supplementary Figure S2**

**Supplementary Figure S2. ESCRT components regulate ubiquitinated cargo sorting in the lymph gland.**

**(A)** Whole-mount larval lymph gland showing accumulation of conjugated ubiquitin (FK2) in the lymph gland upon progenitor-specific (*domeGal4 UAS 2xEGFP* driven) knockdown of 7 *Drosophila* ESCRT components indicated (Vps28, Tsg101, Vps37A, Vps22, Vps32, Vps20, Vps2). Ubiquitin staining is shown in gray scale. Accumulation of ubiquitin aggregates is marked by arrowhead and magnified in insets. Scale bar: 100 µm. **(B)** Bar diagrams show quantification of the fraction of cells accumulating ubiquitin aggregates in primary, secondary and tertiary lobes upon knockdown of all 13 core ESCRT components. n indicates the number of individual lobes analysed and N indicates the number of larvae analysed. Error bars represent SEM. Kruskal Wallis test was performed to determine the statistical significance. *P<0.05, P**<0.01, ***P<0.001. Summary chart indicating presence (colored box) or absence (white box) of ubiquitin accumulation in the primary, secondary and tertiary lobes upon depletion of the respective ESCRT component (left).

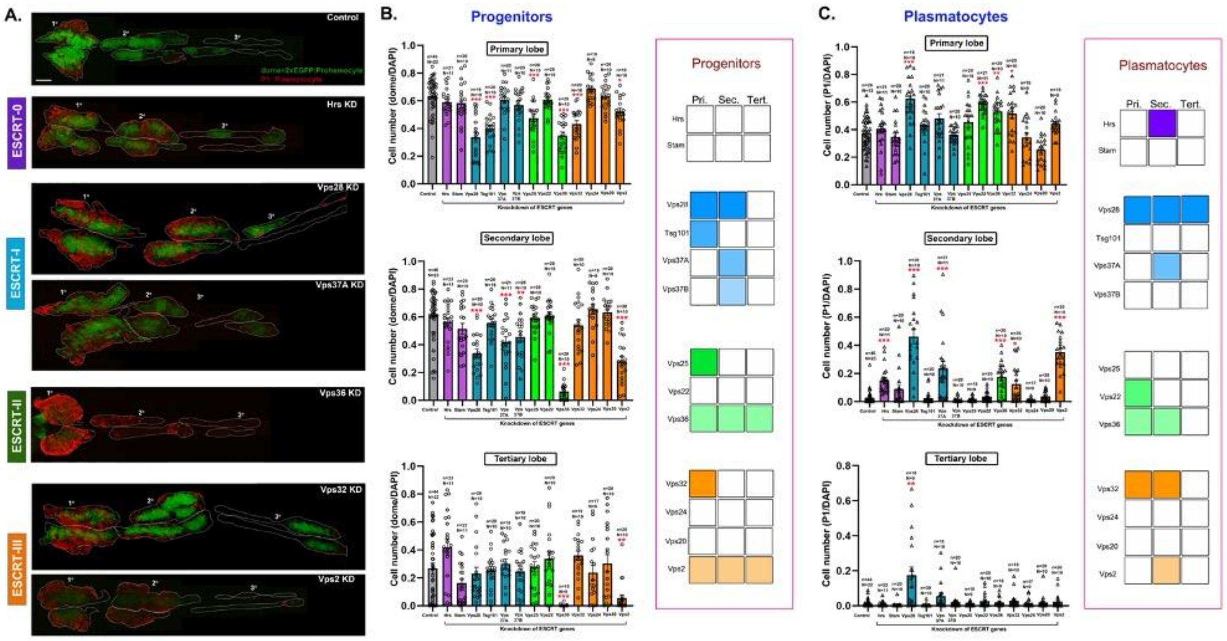


Supplementary Figure S3

**Supplementary Figure S3. ESCRT components regulate progenitor maintenance and plasmatocyte differentiation in the lymph gland.**

**(A)** Whole-mount larval lymph gland showing change in the fraction of dome>2xEGFP+ve progenitors (green) or P1+ve plasmatocytes (red) in the lymph gland upon progenitor-specific knockdown of 6 ESCRT components (Hrs, Vps28, Vps37A, Vps36, Vps32, Vps2). Scale bar: 100 µm. **(B-C)** Bar diagrams show quantification of the fraction of progenitors (B) and plasmatocytes (C) in primary, secondary and tertiary lobes upon knockdown of all 13 core ESCRT components. n indicates the number of individual lobes analysed and N indicates the number of larvae analysed. Error bars represent SEM. Kruskal Wallis test was performed to determine the statistical significance. *P<0.05, **P<0.01, ***P<0.001. Summary chart indicating presence (colored box) or absence (white box) of phenotypes of progenitor loss (B) or increased plasmatocytes differentiation (C) in the primary, secondary and tertiary lobes upon depletion of the respective ESCRT component (left).

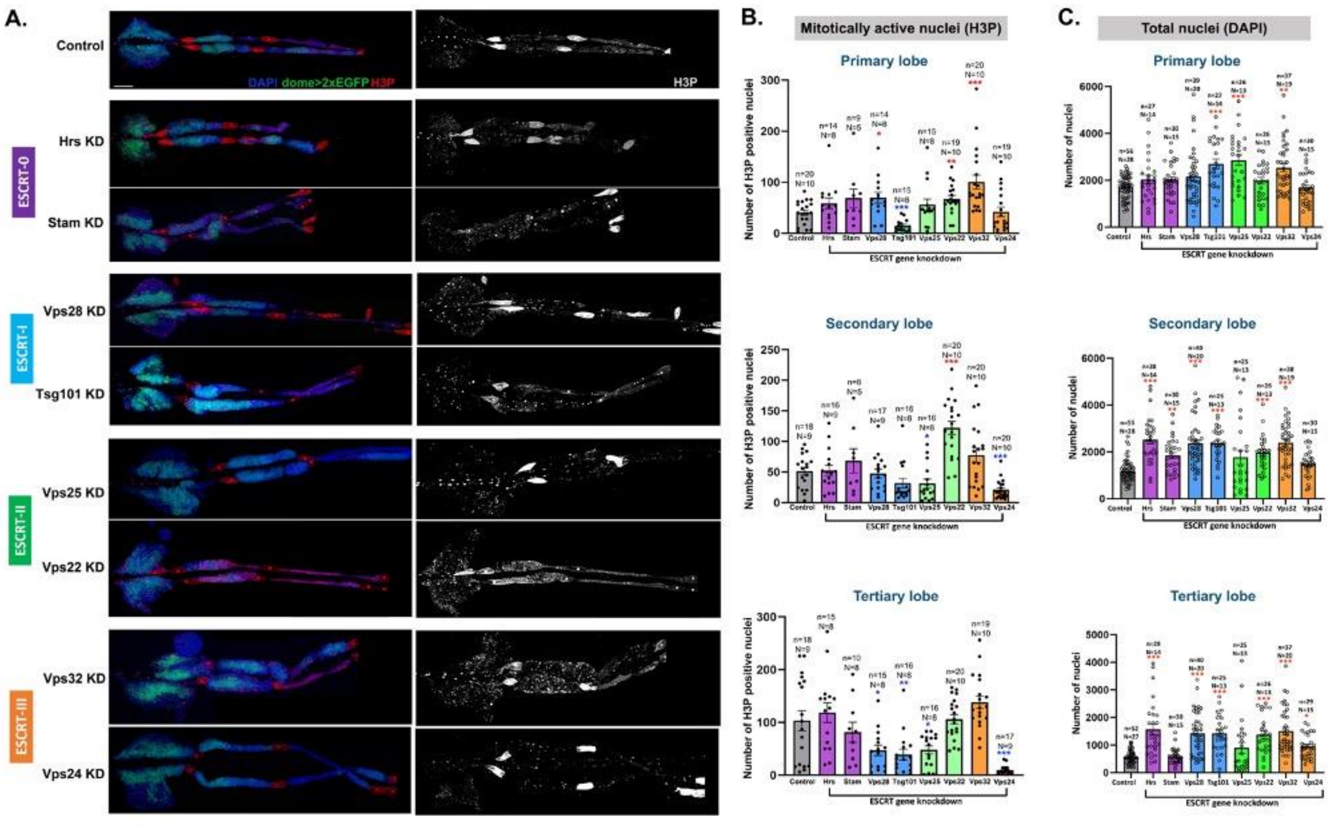


Supplementary Figure S4

**Supplementary Figure S4. ESCRT components differentially regulate mitotic potential and tissue size across the lymph gland.**

**(A)** Whole-mount larval lymph gland showing immunostaining for phosphorylated Histone H3 to mark mitotically active nuclei (red in the left image panel and grayscale in the right image panel) upon progenitor-specific knockdown of 8 ESCRT components [Hrs, Stam (ESCRT-0); Vps28, Tsg101 (ESCRT-I); Vps25, Vps22 (ESCRT-II); Vps32, Vps24 (ESCRT-III)]. Scale bar: 100 µm. **(B)** Bar diagrams show quantification of the number of H3P positive (high H3P) nuclei in primary, secondary and tertiary lobes of the same genotypes. **(C)** The total number of nuclei in each lobe has also been quantified for each lobe in the same genotypes. n indicates the number of individual lobes analysed and N indicates the number of larvae analysed. Error bars represent SEM. One-way ANOVA was performed to determine the statistical significance. *P<0.05, **P<0.01, ***P<0.001.

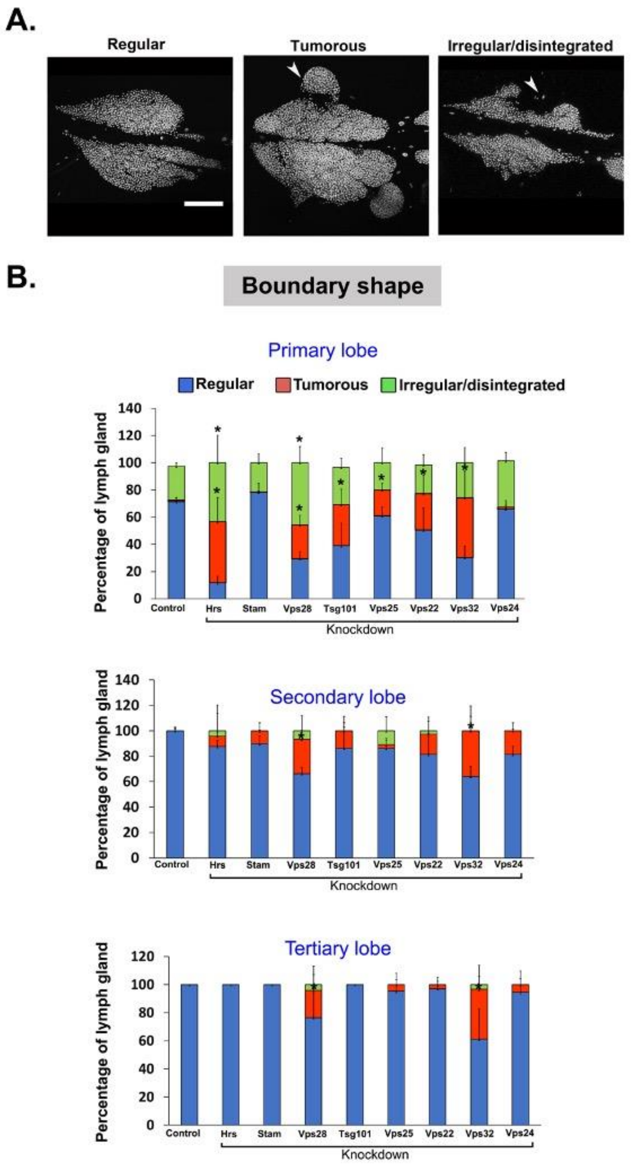


Supplementary Figure S5

**Supplementary Figure S5. ESCRT components affect the morphology of the lymph gland lobes.**

**(A)** Representative primary lobe images of the lymph gland showing regular boundary, irregular/disintegrated boundary and tumorous bulge. Scale bar: 100 µm. **(B)** Quantification of the percentage of larvae showing aforementioned morphology of the primary, secondary and tertiary lobes for knockdown of 8 ESCRT genes [Hrs, Stam (ESCRT-0); Vps28, Tsg101 (ESCRT-I); Vps25, Vps22 (ESCRT-II); Vps32, Vps24 (ESCRT-III)]. N>30 for each genotype. Error bars represent SEM. One-way ANOVA was performed to determine the statistical significance. *P<0.05.

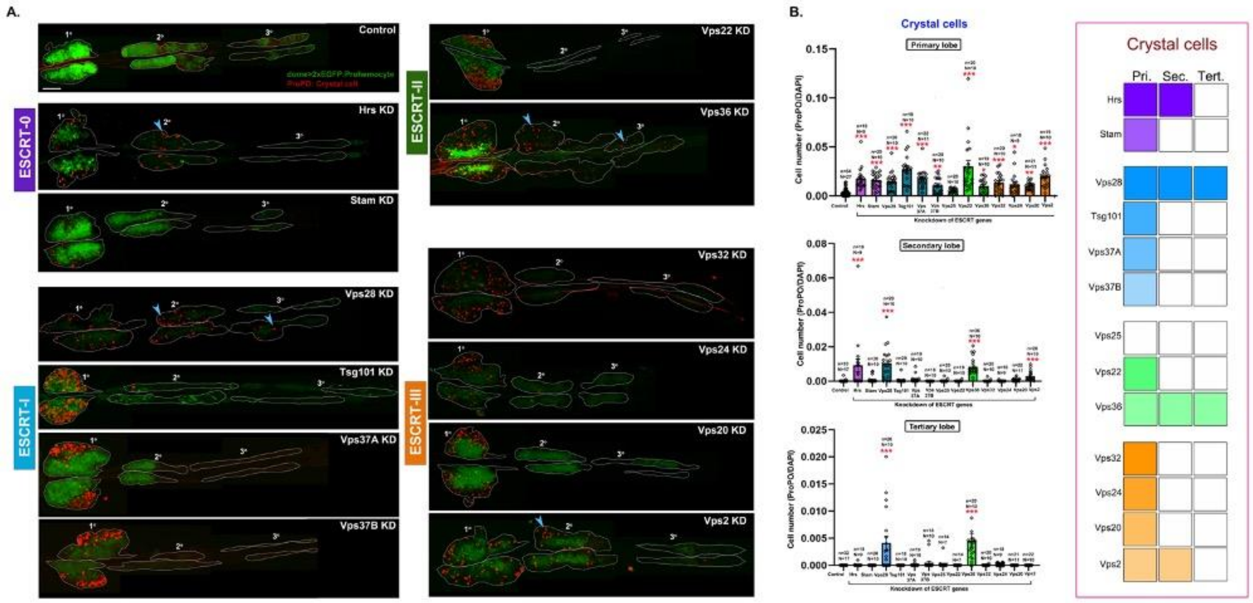


Supplementary Figure S6

**Supplementary Figure S6. ESCRT components differentially regulate crystal cell differentiation of lymph gland progenitors.**

**(A)** Whole-mount larval lymph gland showing differentiation of ProPO+ve crystal cells (red) in the lymph gland upon progenitor-specific knockdown of 12 core ESCRT components. Dome>2xEGFP (green) marks the progenitors across different lobes. Arrowheads mark presence of crystal cells in posterior lobes. Scale bar: 100 µm. **(B)** Bar diagram shows quantification of the fraction of crystal cells in primary, secondary and tertiary lobes upon knockdown of all 13 core ESCRT components. n indicates the number of individual lobes analysed and N indicates the number of larvae analysed. Error bars represent SEM. Kruskal Wallis test was performed to determine the statistical significance. *P<0.05, **P<0.01, ***P<0.001. Summary chart indicating presence (colored box) or absence (white box) of phenotypes of increased crystal cell differentiation in the primary, secondary and tertiary lobes upon depletion of the respective ESCRT component (left). Also see Fig. 1.

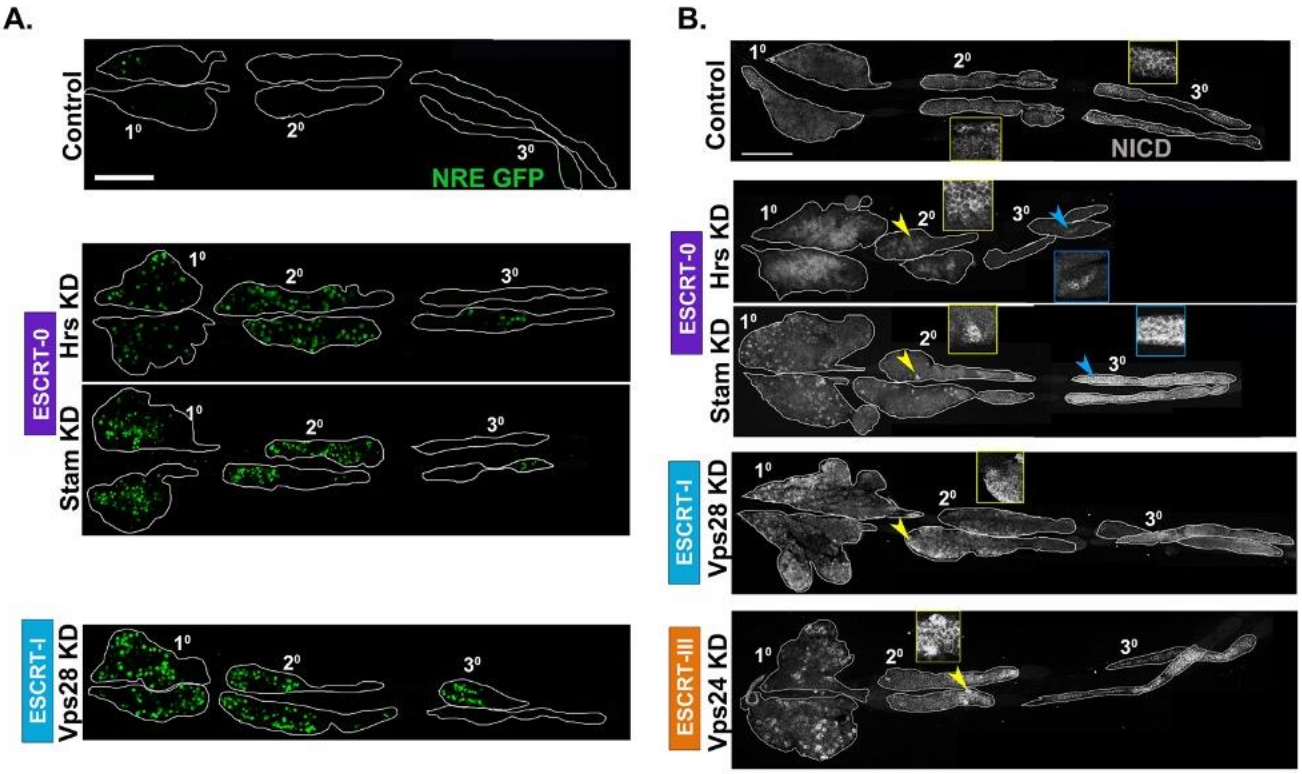


Supplementary Figure S7

**Supplementary Figure S7. ESCRT regulates Notch activation and NICD trafficking in the lymph gland. (related to Figure 2)**

Whole-mount larval lymph gland showing NRE-GFP staining to mark Notch activation across all lobes upon progenitor-specific knockdown of ESCRT components Hrs, Stam (ESCRT-0) and Vps28 (ESCRT-I). Lymph glands in the adjacent panel show NICD staining across the lymph gland upon progenitor-specific knockdown of ESCRT components Hrs, Stam (ESCRT-0) and Vps28 (ESCRT-I). NICD accumulation in secondary (yellow arrowhead) and tertiary lobes (blue arrowhead) are shown in the insets. Scale bar: 100 µm.

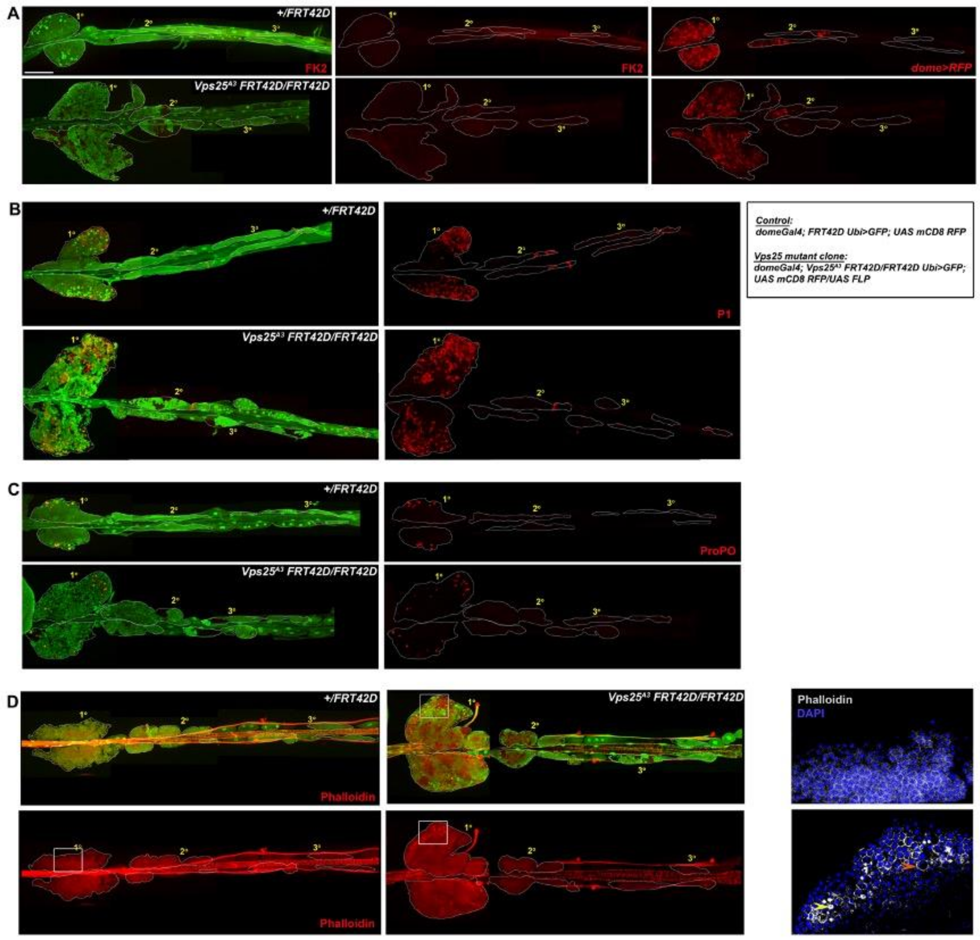


**Supplementary Figure S8**

**Supplementary Figure S8. Vps25 mutation does not affect ubiquitination and blood cell differentiation in the lymph gland. (related to Figure 1, S3, S5, S6 and S9).**

**(A-D)** Whole mount lymph gland showing staining for conjugated ubiquitin in control (*domeGal4; FRT42D Ubi>GFP; UAS mCD8 RFP*) and Vps25 mutant clone (*domeGal4/+; Vps25$^{A3}$FRT42D/FRT42D Ubi>GFP; UAS mCD8 RFP/UAS FLP)* lymph gland. GFP expression marks the wild type twin-spot while GFP negative region marks the homozygous mutant clone. (B) dome>RFP marks the progenitor in the same genotype, (C) P1 marks plasmatocytes, and (D) ProPO marks crystal cells. **(E)** Phalloidin staining was used to visualize lamellocytes based on their elongated morphology. Bottom-most panel shows enlarged view of the boxed region from control and mutant lymph glands. Phalloidin staining in shown in grayscale. DAPI marks the nuclei. The orange arrowhead marks a big binucleate cell while the yellow arrowhead marks a very small cell with high F-actin expression. Scale bar: 100 µm.

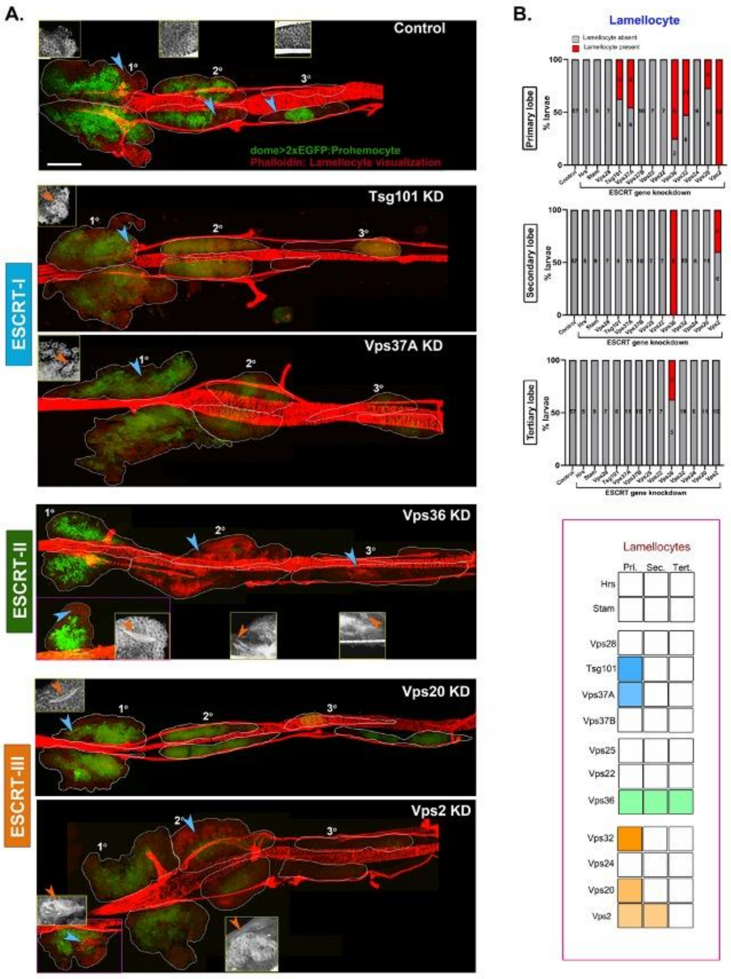


**Supplementary Figure S9**

**Figure S9. ESCRT components regulate lamellocyte differentiation in the lymph gland.**

**(A)** Whole-mount larval lymph gland showing Phalloidin staining (red) to visualise elongated morphology of lamellocytes upon progenitor-specific knockdown of 5 ESCRT components (Tsg101, Vps37A, Vps36, Vps20, Vps2). Blue arrowheads mark the region from primary, secondary or tertiary lobes, magnified in the insets. The inset panel shows enlarged view of Phalloidin staining with lamellocytes marked by orange arrowhead. Scale bar: 100 µm. **(B)** Bar diagram shows quantification of the percentage of lymph glands showing lamellocyte differentiation in primary, secondary and tertiary lobes upon knockdown of all 13 core ESCRT components, without any immune challenge. Values in the columns indicate the number of larvae analysed for presence or absence of lamellocytes. Summary chart indicating presence (colored box) or absence (white box) of lamellocytes differentiation in the primary, secondary and tertiary lobes upon depletion of the respective ESCRT component (left).

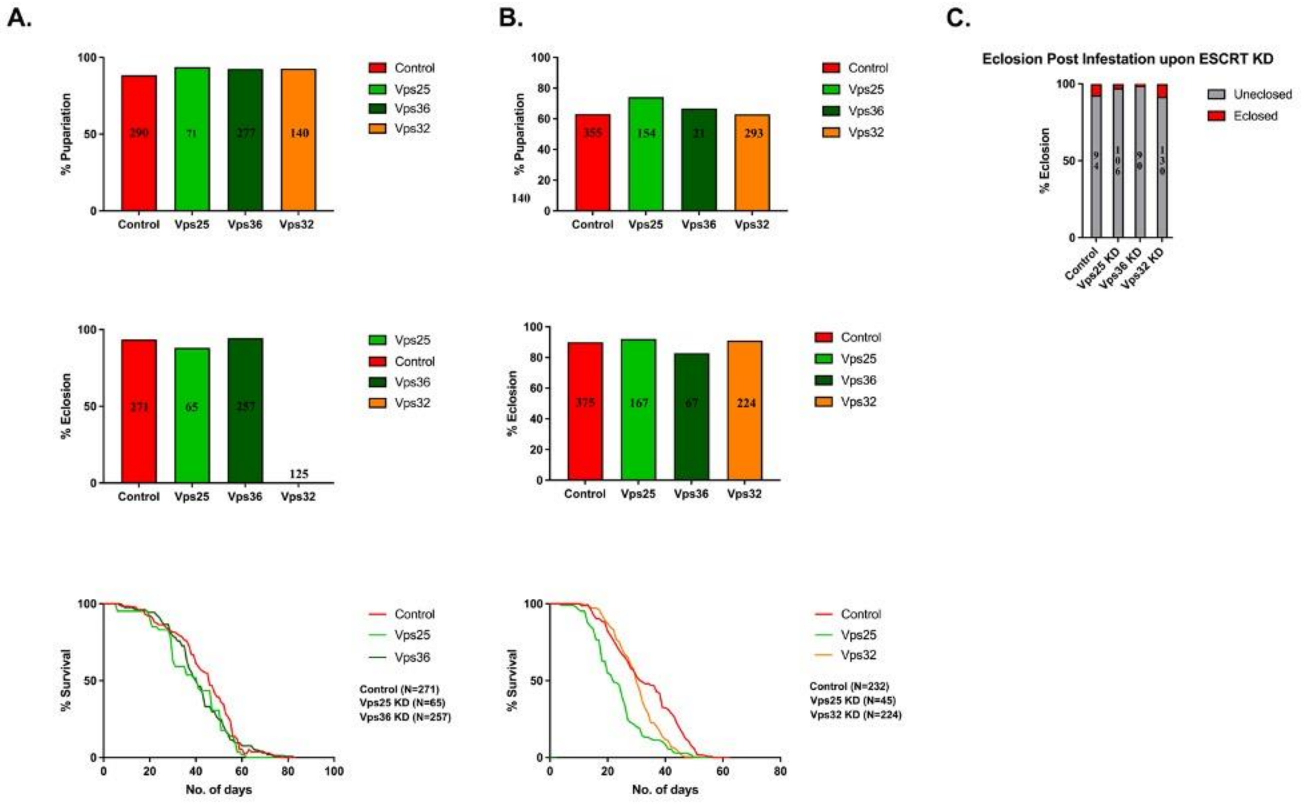


Supplementary Figure S10

**Supplementary Figure S10. ESCRT depletion in hematopoietic compartments does not affect development and survival.**

Developmental analysis of *Drosophila* with Vps25KD, Vps32KD and Vps36KD under **(A)** dome2xEGFP> and **(B)** elav-gal80;; domeMesoGFP> drivers. Quantification in the bar graphs indicate percentage pupariation and eclosion for each genotype. The survival curves visualize percentage of live adult flies per day. **(C)** Quantification visualizes the percentage survival of adult flies post wasp infestation, for flies with Vps25KD, Vps36KD and Vps32KD under elav-gal80 ;; domeMesoGFP> driver. N>90 for each genotype.
